# MassGAT: a graph-based collective learning approach for untargeted detection and annotation of LC-MS data

**DOI:** 10.64898/2026.07.29.741473

**Authors:** Paul-Henri Pinart, Annelaure Damont, Sylvain Dechaumet, Etienne A. Thévenot

## Abstract

**Motivation:** The untargeted processing of Liquid Chromatography coupled to High-Resolution Mass Spectrometry (LC-HRMS) data is a major challenge for the comprehensive and robust characterisation of metabolites. In particular, peak detection and annotation are two challenging tasks due to the size and complexity of the data, that are currently addressed independently in existing pipelines, without taking into account the redundancy of information between the ion species of the same compound. Results: We developed an innovative and efficient approach that combines detection and annotation, by 1) representing signals putatively originating from the same molecule as a graph, and 2) inferring the validity of the peaks and their connections within each component using a Graph Attention Network (GAT). We demonstrate on real data sets that the resulting MassGAT model outperforms current approaches in terms of both detection and annotation. Availability and implementation: The MassGAT open-source Python module is publicly available at https://github.com/odisce/MassGAT.

## Introduction

Metabolomics (i.e., the untargeted characterisation of small molecules ≤ 1.5 kDa in a biological sample) is a major approach for the study of physiopathological mechanisms, and for biomarker discovery (Wishart, 2019; Castelli et al., 2021). High-resolution mass spectrometry coupled with liquid chromatography (LC-HRMS) is the reference technology due to its sensitivity and large dynamic range (Junot et al., 2014; Alseekh et al., 2021). Because of the size and complexity of LC-HRMS data, their treatment (i.e., the characterisation of all metabolites present in the sample) still remains challenging (Guo et al., 2022b; El Abiead et al., 2025).

Raw LC-HRMS data, acquired as scans (mass spectra) at regular time intervals, are recorded as data points in the three dimensions of mass-to-charge ratio (m/z), retention time (rt) and intensity. The signal from an ionized molecule (called a peak) consists of a subset of nearby points forming a local maximum. Each metabolite yields multiple ions species (i.e., multiple peaks) during the electrospray desorption/ionization process (analytical redundancy including isotopes, adducts, fragments, multimers, or multi-charged ions; Nash et al., 2024). Both detection and annotation of the peaks are critical for comprehensive and robust data processing. These tasks, which have been addressed independently so far, are difficult, because of the sparseness and complexity of the signal (including overlap and signal drift), the presence of noise (e.g., contamination by ions present in the solvent, tubing or instrument, as well as electronic noise), and the chemical diversity of analytical redundancy.

On the one hand, numerous signal processing methods have been applied to detect ion peaks (Katajamaa and Orešič, 2007; Renner and Reuschenbach, 2023). Commonly used approaches, available in software such as XCMS (Smith et al., 2006), MZmine (Pluskal et al., 2010), OpenMS (Weisser et al., 2013) or MS-DIAL (Tsugawa et al., 2015), take centroided data as input and proceed in two steps (Renner and Reuschenbach, 2023): 1) search for regions of interest (ROI), e.g., successive data points within a given mass tolerance (centWave: Tautenhahn et al., 2008, FeatureFinderMetabo: Kenar et al., 2014, and ADAP: Myers et al., 2017b), and 2) determination of peaks within these regions (e.g., apex, boundaries and intensity of each peak), using multi-scale approaches based on Continuous Wavelet Transform (centWave: Tautenhahn et al., 2008, and ADAP: Myers et al., 2017b), or the detection of maxima and minima following smoothing (FeatureFinderMetabo: Kenar et al., 2014). These algorithms, however, have certain limitations, such as a significant proportion of false positives and negatives (Myers et al., 2017a; Guo and Huan, 2023), and the complexity of tuning the various parameters (Guo et al., 2022a), which hamper the reproducibility of the analysis of large cohorts, and the adaptability to new biological matrices or experimental setups.

Recently, deep learning methods have been proposed to improve signal detection (Petrick and Shomron, 2022). A few “end-to-end” approaches have been developed that learn both signal detection and quantification directly from the raw data, by using architectures based on YOLO (SeA-M2Net: Zhao et al., 2021) or PointNet++ (3D-MSNet: Wang et al., 2023). However, the labelling of data to train these models is very costly. To simplify the complexity of the input data, alternative approaches thus focus the learning process on pre-identified regions of signal (i.e., peaks or ROIs). Several models have thus been developed to filter Extracted Ion Chromatograms (EIC) from the peaks detected by XCMS, MZmine or MS-DIAL, using a Convolutional Neural Network (CNN) architecture (DNN: Kantz et al., 2019, EVA: Guo et al., 2021, and NeatMS: Gloaguen et al., 2022). Since these approaches do not address the false negatives resulting from the initial peak picking, a second type of methods have been proposed that take ROIs as input rather than peaks, such as the ROIs provided by centWave (peakonly: Melnikov et al., 2019, and QuanFormer: Zhang et al., 2025) or detected as local maxima in the profile data (PeakBot: Bueschl et al., 2022). While demonstrating the value of deep learning for filtering and characterising signals using very few parameters, these models remain sensitive to the initial method used to detect ROIs. In addition, existing methods process peak independently and do not leverage the analytical redundancy to increase the confidence and robustness of the detection.

On the other hand, the grouping of peaks from the same molecule (i.e., the componentization task, as defined by Renner and Reuschenbach, 2023) and their annotation is generally based on a combination of several criteria, including: i) matching against known m/z differences (corresponding to isotopes, adducts, fragments, multi-mers, multi-charged ions, etc.; mz.unity: Mahieu et al., 2016, findMAIN: Jaeger et al., 2017, mzAdan: Stricker et al., 2021, NetID: Chen et al., 2021, and khipu: Li and Zheng, 2023); ii) the similarity of elution profiles (CliqueMS: Senan et al., 2019); and iii) when multiple samples are available, the similarity of intensity profiles (binner: Kachman et al., 2020). These criteria can be combined such as in CAMERA (Kuhl et al., 2012), or supplemented by querying MS1 or MS2 databases when the fragmentation spectrum is available (NetID: Chen et al., 2021). Componentization is critical for subsequent processing steps such as peak alignment between samples (Wandy et al., 2015), for the sensitivity of statistical analyses (Suvitaival et al., 2014) and, ultimately, for the identification of compounds (Böcker et al., 2009; Schmid et al., 2021). However, because current workflows perform annotation as an independent step after the processing is achieved, individual peaks from the same metabolite may be missing or distored as a result of previous smoothing or alignment, thus preventing their correct componentization.

We therefore propose here a paradigm change to tackle the treatment of LC-HRMS data by combining peak detection and componentization, and argue that the grouping of putatively related regions of interest into graph components should enable their efficient selection by deep learning. Our approach (Fig. 1) thus starts with a comprehensive partition of all data points from the raw file by a Watershed-based algorithm, followed by the connection of putatively related ion traces based on known m/z differences and similarity of elution profiles, and, finally, the inference of the true pseudo-spectra and the true peaks by a GNN model. Application of our MassGAT module on several real data sets from various instrumental settings shows the value of this new collective learning approach for LC-HRMS data processing and annotation in metabolomics.

**Figure 1.**
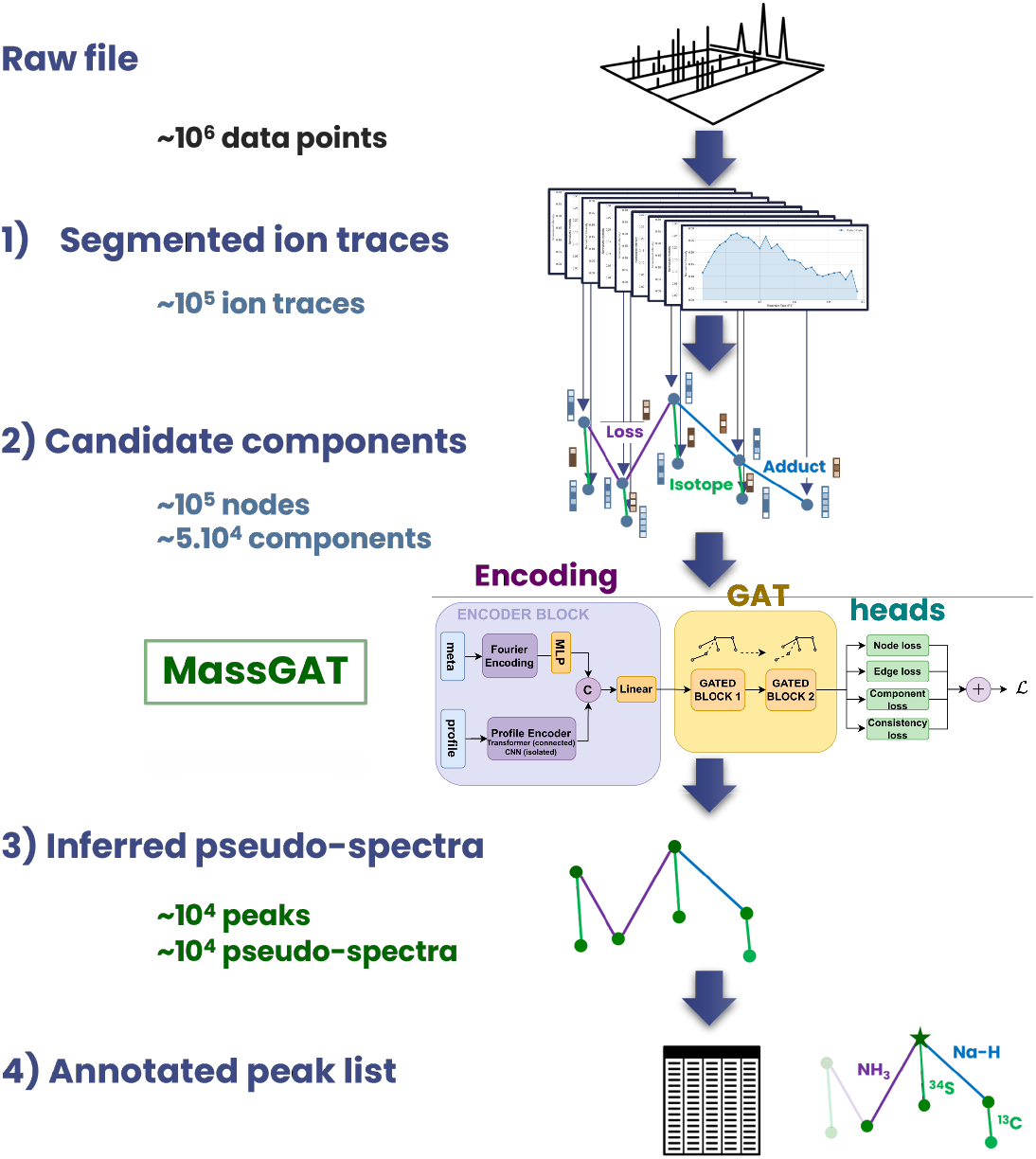
Overview of the LC-HRMS combined data processing and annotation by MassGAT. After comprehensive segmentation of all data points from the raw file (1), segments (called ion traces) are connected in candidate components based on m/z differences and similarity of elution profiles (2). The MassGAT GNN then prunes these graphs to discard noise components and noise segments (3). The resulting components with validated peaks only (called pseudo-spectra) are annotated and the final peak list is returned (4). The order of magnitude of the number of signals of interest is indicated at each step.

### Approach

We have developed a novel joint inference framework for untargeted LC-HRMS preprocessing and annotation (Fig. 1). Our software takes as input a centroided raw file (.mzML format) and returns the list of detected peaks, with their annotations and visualizations. It consists of four mains steps: 1) segmentation of all *data points* (i.e., triplets of mass-to-charge ratio, retention time, and intensity) into sets corresponding to local maxima, that are referred to as *ion traces*; 2) grouping ion traces putatively related to the same metabolite as an undirected graph called *component* based on chemical rules (co-elution, similarity of their elution profiles, and matching of *m/z* differences to known chemical relationships); 3) processing each component by a Graph Attention Network (GAT; Fig. 2) to infer which nodes (ion traces) are true peaks and which edges are true chemical relationships; 4) annotating the resulting *pseudo-spectrum* by characterizing the main ion and determining the neutral mass of the compound (Fig. 3).

**Figure 2.**
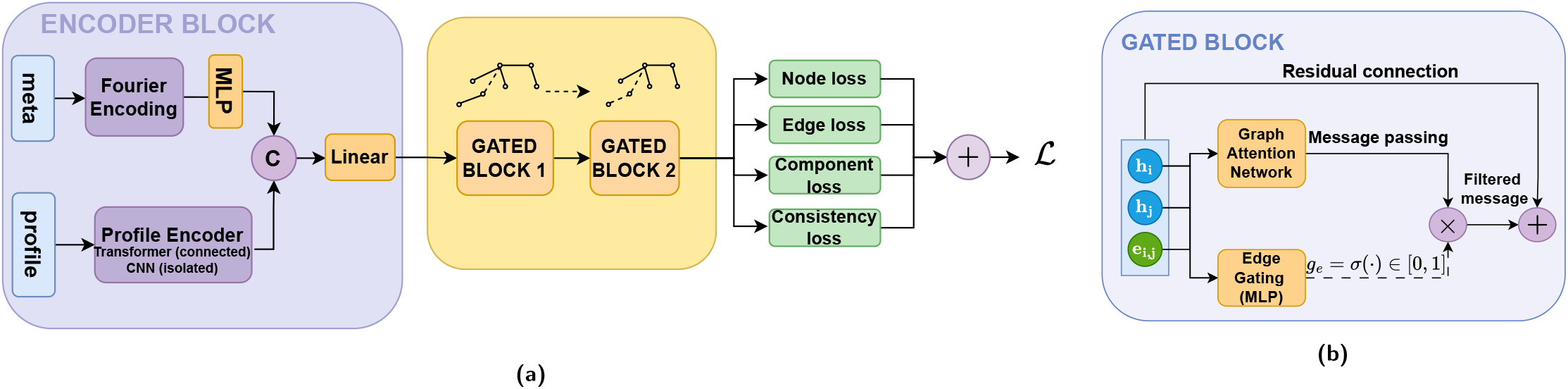
Architecture of the MassGAT neural network. **(a)** Overview of the full model. The feature vector of each ion trace is split into scalar meta-features and a chromatographic profile, encoded separately and concatenated before a linear projection gives the initial node representation 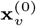. Connected ion traces are processed through a Transformer profile encoder, followed by two successive gated blocks to refine node representations through message passing. Isolated ion traces are processed through a CNN encoder only. Four loss terms are computed from the final embeddings and combined into the training objective (Equation 2). **(b)** Detail of a single gated block. Node representations *h*_*i*_, *h*_*j*_ and edge attribute *e*_*ij*_ are passed in parallel to a Graph Attention Network and to an edge-gating MLP. The scalar gate *g*_*e*_ = *σ*(*·*) *∈* [0, 1] scales the aggregated message before it is added back via a residual connection. Each gated block contains *L* = 2 such layers.

**Figure 3.**
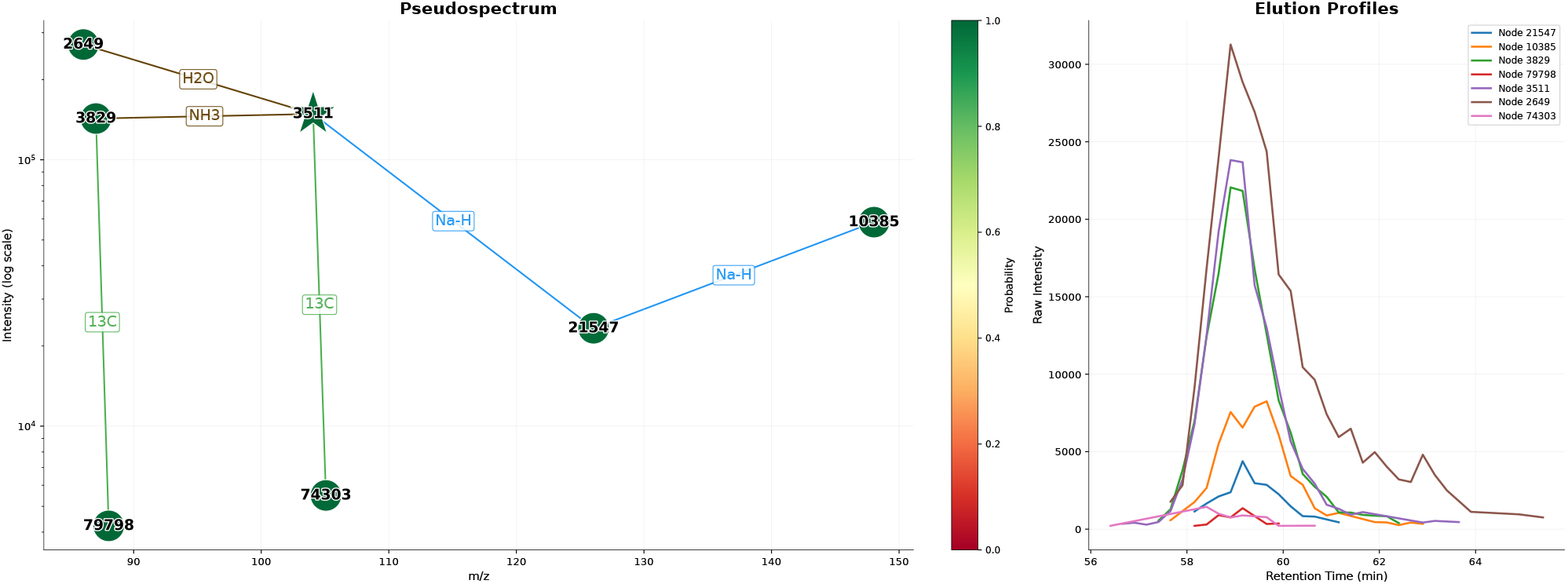
Visualization of the pseudo-spectrum (i.e., component validated by MassGAT) corresponding to GABA in the IRS2-KO sample (positive mode). **Left:** The annotated pseudo-spectrum is shown in the *m/z* vs. intensity space (left). The star corresponds to the targeted ground-truth peak. Nodes (detected ions) are coloured according to their detection probability (green: 1, yellow: 0.5). Edge are labeled by the chemical formula of the m/z difference, and coloured according to type of link (green: isotope, blue: adduct, purple: loss, brown: loss or neutral adduct). The base peak annotated as [M+H]^+^ is observed at *m/z* = 104.0696, and associated with isotopic (^13^C), adduct (Na-H) and loss links (H_2_O and NH_3_). **Right:** Elution profiles from the nodes (i.e., peaks) of the pseudo-spectrum.

Our main contributions are: (i) the formulation of peak detection and componentization as a simultaneous node, edge, and graph classification task on a chemical graph; (ii) a hybrid encoding architecture that applies Transformer and GNNs to connected nodes to capture global coherence, and CNNs to isolated singletons to detect local shape quality; (iii) a semi-supervised loss function that enables to include partially annotated real components as training data without penalizing the model; (iv) a robust synthetic data generation pipeline that samples peak parameters and chemical relationships to augment the pool of graph components for training; and (v) a more consistent processing software for LC-MS data, including interactive visualizations to facilitate the interpretation of pseudo-spectra.

## Materials and Methods

### Segmenting the data points into ion traces

The centroided raw file (provided in open format such as mzML) is segmented (i.e., each recorded data point [m/z, rt, intensity] is assigned to a segment, called an ion trace) by using a Watershed algorithm on both m/z and rt dimensions (https://github.com/odisce/ionshed). This methodology ensures that most segments contain at most one true ion peak. Note that alternative approaches to extract ion chromatograms can also be used at this step (Renner and Reuschenbach, 2023). Segments with less than 4 data points (user parameter) were discarded as containing not enough information to discriminate between signal and noise. As an example, the segmentation step typically generates 4 *×* 10^5^ ion traces containing on average 8 data points [min = 4, max = 99] (respectively, 29 data points [min = 4, max = 620]) when applied to a sample from the TripleTOF 6600 (respectively, QE HF) dataset.

### Representing putatively related ion traces as a graph component

Each segmented ion trace (i.e., a list of data points) is represented as a 72-dimensional feature vector, with 8 meta features to characterize the segment (center coordinates, integrated intensity), and 64 features to include the elution profile. The original profile is either padded by zeros or interpolated to match this fixed dimension. The retention times are shifted to [0, rt_max_ *−* rt_min_] and the intensities are normalized by the maximum value.

To leverage the structural dependencies between ions of the same metabolite, the segments are connected within an undirected graph where edges correspond to chemical links such as isotopes, adducts, fragments, or dimers (see the default list of m/z differences in Table S1). This graph is built hierarchically by connecting isotopes, then adducts and fragments, and finally dimers (Note S1).

### Processing each component by a GAT architecture

They key idea of our collective learning approach is that the presence of neighbours within a component at known m/z differences and with correlated elution profiles should reinforce their probability of being true peaks, while, conversely, the model should penalize the absence of any connected nodes with a similar shape. We therefore propose a GNN architecture where each node iteratively aggregates information from its chemical neighbours, so that the final classification of each ion reflects both its own shape and the consistency of its chemical context (Fig. 2). The model simultaneously yields three scores: at the node level (peak vs. noise), at the edge level (truly related ion species vs. accidental connection), and at the component level (pattern from a real metabolite vs. component containing only noise). The architecture of the MassGAT model (Fig. 2) and its implementation are detailed in the Note S1.

Briefly, node encoding is performed using random Fourier features followed by a Multilayer Perceptron (MLP) for the meta features, and using a Transformer (respectively, a CNN) for the elution profiles of the connected (respectively, isolated) nodes. Edge attributes encode the type of chemical relationship and metrics about the respective intensities, retention times, and correlation of elution profiles between the adjacent nodes.

Initial embeddings are passed through graph attention layers (Veličkovićet al., 2017) with a learned edge-gating mechanism (Bresson and Laurent, 2017), *g*_*e*_ *∈* [0, 1], computed for every edge *e* = (*u, v*):

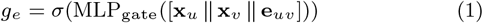

where *σ* is the sigmoid function. The gate prevents aggregation from noisy nodes into a clean component. After the final block and the shared projection layer, the three projection heads are run in parallel. The model contains 98,000 trainable parameters.

The loss combines four terms:

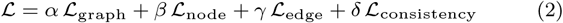

The consistency loss enforces agreement between each edge and its connected nodes, as well as between the component and its most probable node (Note S1).

### Augmenting the training data with simulated components

A total of 18,000 real data were used for training, including about 3,000 partially annotated true components from an in-house database (Roux et al., 2012; Boudah et al., 2014), as well as 10,000 noise components and 5,000 noise isolated ion traces that were extracted from a blank sample with a zigzag index metric above 0.2 to ensure that only real negatives were kept (Zhang et al., 2023). Since the real components are partially annotated, a semi-supervised setting was implemented in MassGAT so that the predictions for the nodes that have not been manually annotated are excluded from the MassGAT losses (Note S1).

These data were augmented with 70,000 synthetic data (Note S1), including 40,000 simulated components (20,000 components with noise only, 10,000 components with true peaks only, and 10,000 components mixing both types of ion traces) in addition to 20,000 synthetic isolated noise traces and 10,000 synthetic isolated true peaks. Elution profiles were simulated with Exponentially Modified Gaussians for the true peaks (Grushka, 1972) or random noise, oscillation or drifts for the noise ion traces. Realistic measurement noise was added in both cases. Components were generated by selecting a main ion and then deriving its isotopic cluster, adducts, fragments and dimers, by using true peaks or noise ion traces with high cosine similarity and matching m/z differences. Expected natural abundance ratios of intensities were used for the isotope peaks.

### Annotating the validated pseudo-spectra

The pseudo-spectra are annotated by 1) assigning the main ion (by default the (de)protonated molecule), 2) propagating the chemical relationships in a directed manner, and 3) computing the neutral mass of the metabolite, as described in Note S1 and summarized below. Candidate main ions (i.e., peaks not already annotated as isotope or dimer child) are assigned a score based on the normalized number of children, isotope children, predicted probability and intensity (Note S1). The heuristic relies on the observation that the true main ion usually acts as a root, with isotope, adducts and dimers pointing back to this node. In contrast to the initial component which is built as an undirected graph, here, for each candidate main ion ([M+H]^+^ or [M-H]^*−*^ by default), its edges can be assigned a direction (i.e., adducts must propagate towards heavier nodes, and losses towards lighter nodes), enabling to compute a global annotation score. A softmax distribution is finally computed to give a probabilistic estimation of the best candidates. The neutral mass of the compound is estimated by weighted regression over all non-dimer peaks. The five top scoring annotations are included in the peak list. The pseudo-spectra, with the annotations and the elution profiles can be visualized interactively (Fig. 3).

### Test datasets

Two sets of real data were used for the benchmark of the peak detection and annotation tasks, respectively.

#### TripleTOF 6600 and QE HF

Two mixtures of 1,100 compounds, spiked at specific concentrations in black pepper extracts, have been analyzed in quadruplicate on a C18 column coupled to either an AB SCIEX TripleTOF 6600 or a Thermo Q Exactive HF instruments in the positive ionisation mode (Li et al., 2018). Using a targeted data analysis with the vendor software, peaks from 970 and 836 compounds have been identified by the authors in the TripleTOF 6600 and QE HF data, respectively, and thus define the ground truth. The raw data are publicly available.

#### IRS2-KO and MTBLS103

The Retina IRS2 data consist of the LC-MS1 acquisition from retina samples of Irs-2-deficient mice on a C18 column coupled to a QTOF instrument in both ionization modes (Senan et al., 2019), and includes the peaks from 20 metabolites manually confirmed by MS2 fragmentation. The MTBLS103 dataset comprises serum samples from a cohort study of HyperInsulinaemic Androgen Excess (HIAE), that were analysed in the positive mode on a QTOF instrument with two chromatographic conditions: 18 samples on a C18 column and 13 samples on a HILIC column, with 9 and 6 confirmed metabolites respectively (Samino et al., 2015). The raw data from the Retina IRS2-KO and the HIAE study are publicly available at zenodo.1480659 and MTBLS103, respectively. The processed and annotated data from the previous benchmark on these data (Senan et al., 2019) were kindly provided by the authors.

## Results

We have developed a new model for the processing LC-HRMS data by chemistry-informed deep learning which, based on the exhaustive partition of the raw data, connects the segments potentially originating from the same metabolite into graph components, and infers the true pseudo-spectra and the true peaks with a gated GNN (Fig. 1). The MassGAT model was successfully trained on a mixture of real and simulated components, achieving an accuracy of 99% and 98% at the node and graph levels, respectively (Note S2).

### Peak detection

The quality of peak detection by MassGAT was assessed on the TripleTOF 6600 and QE HF Orbitrap benchmark datasets (spiked black pepper samples acquired in quadruplicate on both instruments; Li et al., 2018): MassGAT was first applied to each file and the detected peaks were subsequently aligned with XCMS (Smith et al., 2006) to generate the final peak table (Note S1).

All files were efficiently processed by MassGAT (Note S1), with running times for the whole process (segmentation, building of the components, inference of the true pseudo-spectra and peaks, and annotation) of 2.3 min for a TripleTOF 6600 file (respectively, 8.4 min for a QE HF file) on 5 CPU cores, without requiring GPU hardware. A critical aspect of the MassGAT is its massive pruning of the initial ion trace candidates, the vast majority of which represent electronic or chemical noise: the acceptance rate was 19.2% in the TripleTOF 6600 files for isolated singletons (respectively, 11.6% in the QE HF files), and 45.5% (respectively, 21.7%) for ion traces within graph components (Fig. S1). The average number of peaks per pseudo-spectrum was 2.3 and 2.5 in the TripleTOF 6600 and QE HF samples, respectively (Fig. S2). These results indicate that while the model values the chemical context, it still performs active filtering within valid components.

MassGAT provided the best recalls compared to the alternative software XCMS (Tautenhahn et al., 2008), asari (Li et al., 2023), and 3D-MSNet (Wang et al., 2023), achieving 97.3% and 99.2%, respectively (Table 1). The comparison of the targets detected the MassGAT, XCMS and 3D-MSNet software is shown on Fig. S3. Importantly, MassGAT was also shown to outperform the CNN approach (which filters ion traces individually, without considering potential chemical relationships), highlighting the value of the collective learning approach (Table 1).

**Table 1.** Peak detection performance on the TripleTOF 6600 and QE HF datasets. Consensus true features are recovered in all files with a probability threshold of 0.5 for at least 50% of replicates (≥ 4/8). The true feature ID rate is the percentage of identified ground-truth features. Total features refers to all ion traces retained at probability ≥ 0.5 (MassGAT and CNN) or by the XCMS, asari and 3D-MSNet software.

| Method | Consensus true features | True feature ID rate (%) | Total features |
| --- | --- | --- | --- |
| *TripleTOF 6600 (970 targeted truth features)* |  |  |  |
| XCMS | 892 | 92.0 | 24 088 |
| asari | 783 | 80.7 | 65 376 |
| 3D-MSNet | 939 | 96.8 | 22 526 |
| CNN | 932 | 96.1 | 40 324 |
| MassGAT (ours) | **944** | **97.3** | 32 566 |
| *QE HF Orbitrap (836 targeted truth features)* |  |  |  |
| XCMS | 823 | 98.4 | 25 702 |
| asari | 355 | 42.5 | 26 699 |
| 3D-MSNet | 828 | 98.9 | 40 386 |
| CNN | 827 | 98.7 | 32 874 |
| MassGAT (ours) | **829** | **99.2** | 39 742 |

One of the objectives of our collective learning approach is that it should help detecting valid isotopes, adducts and dimers that can be of low-intensity or irregular shape, if they form a coherent component with more standard ion traces. We indeed observed that isotopes of low intensity but consistent intensity ratios were detected by MassGAT (Fig. S4 and Fig. S5).

To further test this hypothesis, we evaluated the presence of the ^13^C isotope for each recovered ground-truth compound: the theoretical mass and expected abundance of the ^13^C isotope were calculated and searched in the consensus peak tables (*±*10 ppm for *m/z, ±*0.3 min for retention time, and a 30% abundance tolerance). MassGAT identified the highest number of valid isotopic pairs on both datasets (Fig. 4). On the TripleTOF data, MassGAT validated 719 isotopes out of 944 recovered features (76.2%), compared to 643/892 (72.1%) for XCMS and 561/939 (59.7%) for 3D-MSNet. On the QE HF Orbitrap dataset, a high validation rate was also obtained by MassGAT (721/829, 87.0%), matching XCMS (715/823, 86.9%) and outperforming 3D-MSNet (639/828, 77.2%).

**Figure 4.**
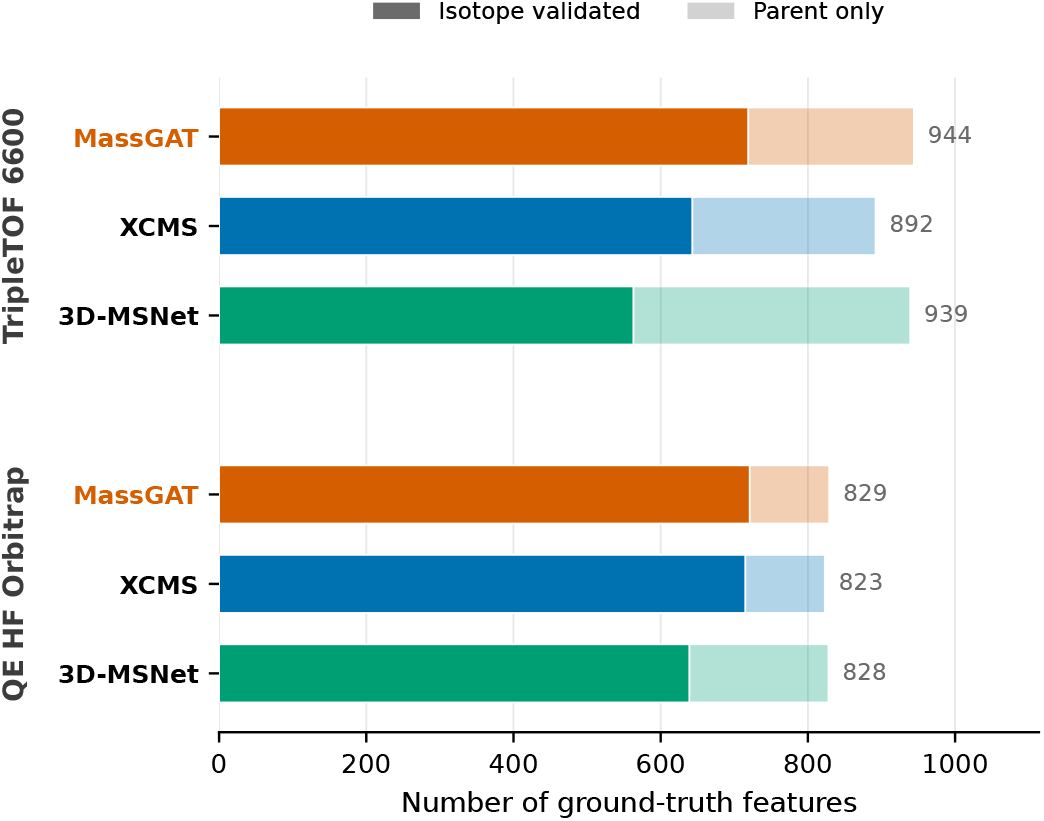
Detected ground-truth features along with their ^13^C isotope across the TripleTOF 6600 and QE HF Orbitrap datasets. Dark (respectively, light) colors indicate ground truth features for which the expected ^13^C isotope was successfully validated within abundance tolerances (respectively, features where only the parent ion was recovered).

### Componentization and annotation

Componentization and annotation were assessed on the Retina IRS2-KO and MTBLS103 datasets, previously used by Senan et al., 2019 to benchmark the performances of their CliqueMS software (which, as MassGAT, is designed to process files individually), and the alternative approaches CAMERA (Kuhl et al., 2012), xMSannotator (Uppal et al., 2017), and MS-FLO (DeFelice et al., 2017). As a baseline, CliqueMS was rerun to include the additional annotated m/z differences from MassGAT.

In the retina IRS2-KO samples, MassGAT correctly ranked the true neutral mass among the top 5 candidates for all ground-truth metabolites in the positive ionization mode, outperforming the 16 out of 20 correct annotations obtained by CliqueMS (Table 2 and Table S3). In the negative mode, 16 metabolites (out of 18) were annotated by MassGAT, compared to 6 for CliqueMS (Table 2 and Table S4). Note that, to increase the confidence in the annotation results, only metabolites with at least two detected ion species (i.e., with two nodes in the pseudo-spectrum) were considered as annotated. All types of ion species were enriched in most MassGAT pseudo-spectra (Fig. 5). In particular, some species that were entirely undetected by CliqueMS (e.g., L-Glutamic acid and Taurine in negative mode), or with a poor network (L-Ascorbic acid in positive mode), were successfully recovered by MassGAT, providing a more exhaustive characterization of the mass spectrum.

**Table 2.**
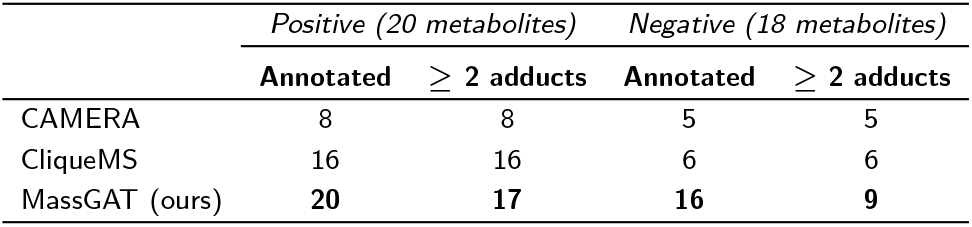
Annotation of the retina IRS2-KO samples. The performances for CAMERA are those reported from the previous benchmark (Senan et al., 2019).

**Figure 5.**
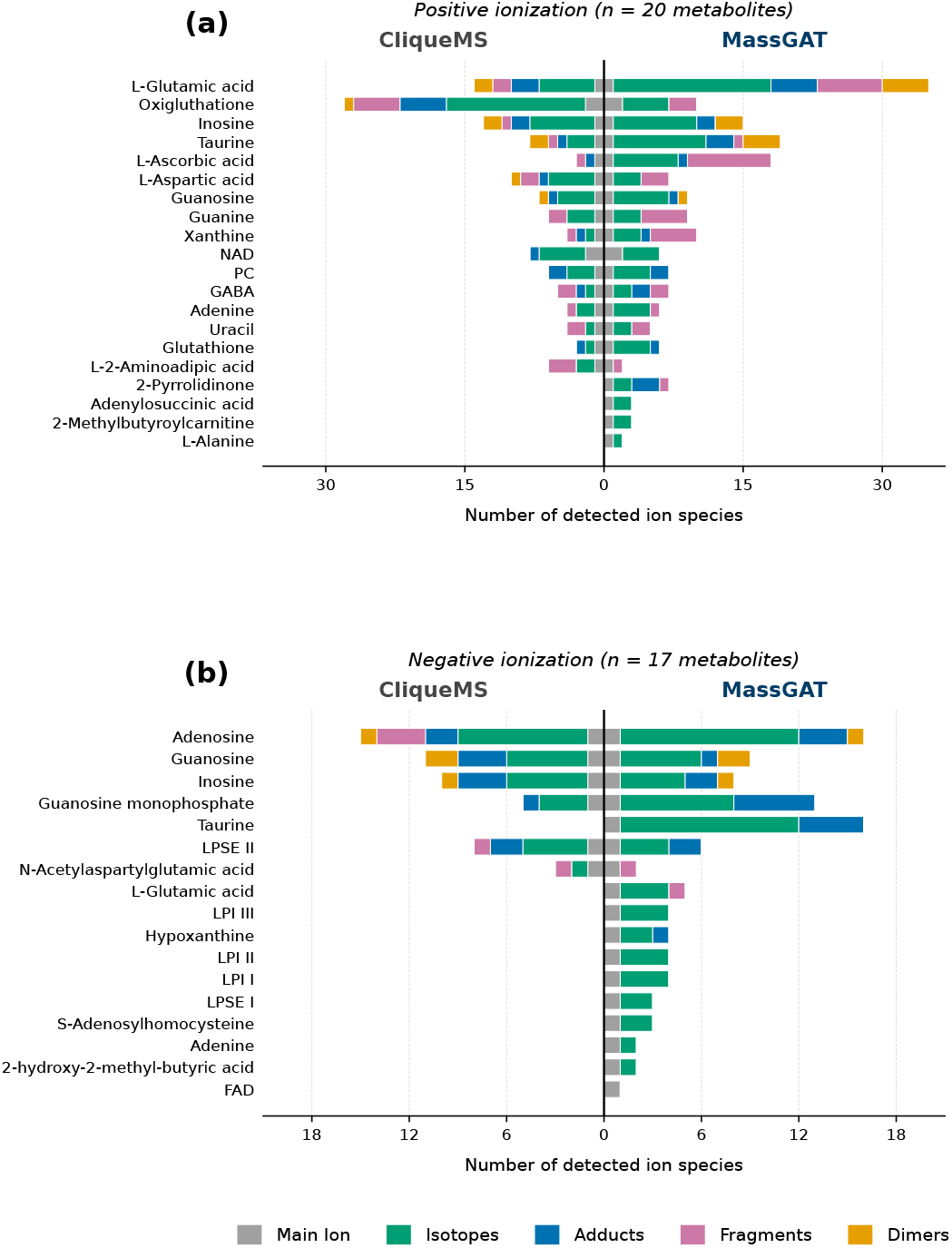
Annotated ion species in the IRS2-KO samples from the positive (**a**; *n* = 20 ground truth metabolites) and negative (**b**; *n* = 17 metabolites) ionization mode.

Annotations can be displayed interactively within the pseudo-spectrum along with the elution profiles for manual exploration (Fig. 3, Fig. S4 and Fig. S5). Interestingly, MassGAT automatically detects all sequences of the same chemical relationship (e.g., two successive Na-H adduct links in GABA and multiple Na + Cl neutral adducts in Taurine; Fig. 3 and Fig. S5, respectively), in contrast to existing software where all links to be searched have to be explicitly listed in the input table of m/z differences. Furthermore, in two cases, MassGAT provided new identification information compared to the original annotation by Senan et al., 2019. For GABA (Fig. 3), a loss of H_2_O matches the mass of the protonated 2-Pyrrolidinone (*m/z* = 86.0595). Since both ions have the same elution profiles (brown and purple curves, respectively), we would assume that the ion *m/z* = 86.0595 is an in-source fragment of GABA and not the protonated ion of 2-Pyrrolidinone. For L-Aspartic acid, the pseudo-spectrum (Fig. S6) suggests that the main ion *m/z* = 134.0434 is in fact an in-source fragment resulting from a C_2_H_2_O loss (see the perfect alignement of the elution profiles in red and brown): consequently, the observed metabolite is more likely N-Acetylaspartic acid than L-Aspartic acid.

For the multi-sample HIAE cohort (MTBLS103), annotation consistency was evaluated across all biological replicates using three standardized metrics: *r*_1_ (correct annotation in the top 5 in *>* 50% of the samples containing the target), *r*_2_ (correct annotation in at least one sample), and *r*_3_ (the total cumulative hits across the entire cohort). MassGAT demonstrated reproducibility across both the C18 and HILIC chromatographic conditions (Table 3, Table S5, and Fig. S7).

**Table 3.** Annotation of the samples from the HIAE cohort (MTBLS103 dataset). *r*_1_: correct in *>* 50% of samples; *r*_2_: correct in ≥ 1 sample; *r*_3_: aggregate over all samples. The performances for xMSannotator, MS-FLO, and CAMERA are those reported from the previous benchmark by Senan et al. 2019.

| Method | $r_1$ ($> 50\%$) | $r_2$ ($\geq 1$) | $r_3$ (Total) |
| --- | --- | --- | --- |
| *C18 (18 samples, 9 metabolites)* |  |  |  |
| xMSannotator | 3 | — | — |
| MS-FLO | 0 | — | — |
| CAMERA | 3 | — | — |
| CliqueMS | 6 | 8 | 104 |
| MassGAT (ours) | **8** | **9** | **133** |
| *HILIC (13 samples, 6 metabolites)* |  |  |  |
| xMSannotator | 1 | — | — |
| MS-FLO | 1 | — | — |
| CAMERA | 3 | — | — |
| CliqueMS | **5** | **6** | **56** |
| MassGAT (ours) | **5** | **6** | **72** |

For the *r*_1_ metric, MassGAT correctly annotated 8 out of 9 (respectively, 5 out of 6) metabolites acquired in the C18 (respectively, HILIC) conditions in the majority of the replicates (Table 3). In contrast, CliqueMS annotated 6 metabolites out of 9 in the C18 data. Furthermore, the aggregated annotations (*r*_3_) show the robustness of the collective learning framework across variable sample conditions, with 133 and 72 annotations (vs. 104 and 56 with CliqueMS) for the C18 and HILIC data.

## Discussion

We have developed a new approach to process LC-HRMS data that simultaneously detects and annotates peak patterns of ion species from the same metabolite. Using real datasets obtained with two distinct instruments (TOF and Orbitrap), two chromatographic columns (C18 and HILIC), and both ionisation modes, we show that the number of target compounds detected is higher than that achieved by conventional approaches such as XCMS/centWave (Tautenhahn et al., 2008) and asari (Li et al., 2023), as well as a recent deep learning method based on point cloud segmentation (3D-MSNet: Wang et al., 2023). We also show that the number of annotated peaks exceeds that of alternative approaches such as CliqueMS (Senan et al., 2019) and CAMERA (Kuhl et al., 2012). These results demonstrate the value of combining peak detection and componentization, which, to our knowledge, had never been proposed before: the simultaneous detection of peaks within a group enhances prediction accuracy (as shown by MassGAT’s superior performance compared to a CNN filter applied to individual ion traces).

To process peaks collectively, MassGAT relies on modelling ion patterns as graphs. Trees or graphs have been used to represent information about the relationships between peaks in MS1 (Kuhl et al., 2012; Mahieu et al., 2016; Li and Zheng, 2023) and MS2 spectra (Rasche et al., 2011; Delabri`ere et al., 2025). Recently, a graph representation of MS1 imaging data has been used to learn the spatial segmentation with a relational graph convolutional network (La Rocca et al., 2025). In MassGAT, the representation is enriched to incorporate information from elution profiles into the nodes (in addition to the relationships between ion species in the edges) and therefore to enable reasoning about patterns based on all available information through a GAT architecture.

MassGAT is designed for routine use in untargeted metabolomics analysis. On the one hand, its computation times (in the order of a few minutes per sample) are comparable to those of existing software. On the other hand, MassGAT integrates with downstream modules (such as those from XCMS) for alignment between samples. Finally, the lightweight architecture of the model allows for retraining even on CPU resources, and the code repository adheres to good practices in machine learning to facilitate its reuse.

The main constraint of the current model is that the components (and therefore the pseudo-spectra) rely on a list of m/z differences provided in the parameters. Consequently, MassGAT may return several pseudo-spectra for the same compound if the components are linked by an m/z difference that is not present in the list. To minimise this risk, the default list has been selected from m/z differences frequently encountered in the data, and the user is free to modify it to suit their experimental setup (e.g., specific salts present in the mobile phases). To take this a step further, a post-processing step could be added to indicate to the user which pseudo-spectra are likely to originate from the same compound.

The model is trained on both real and synthetic data, in order to ensure a sufficient amount of realistic data. Opting for a fully parametric simulation of elution profiles, rather than augmenting empirical true peaks, makes it possible to compensate for the lack of perfectly isolated reference peaks in real-world datasets. Furthermore, this strategy ensures a deeper and controlled exploration of the analytical space by generating a wide variety of potential peak shapes and boundary scenarios. The detection accuracies on both Time-Of-Flight and Orbitrap instruments demonstrate that the proposed simulator is realistic. Recently, an LC-HRMS data file generator was described in the mzrtsim package (Yu and Philip, 2025). Our initial comparisons suggest that the diversity of peak shapes and noise generated by MassGAT more closely resembles real data. However, the mzrtsim method has the advantage of using real spectra to construct peak patterns, which appears to be a promising approach for further enhancing the realism of the data simulated by MassGAT. In conclusion, MassGAT is an innovative and effective approach for chemistry-informed learning of the detection and annotation of LC-HRMS data, which are critical for the discovery of metabolite biomarkers. The ability to detect and utilise analytical redundancy directly within the raw signals opens up new possibilities for improving the sensitivity and robustness of all subsequent steps of metabolomic data processing (e.g., alignment between samples), statistical analysis, and annotation (e.g., confidence levels).

## Acknowledgements

We thank Thomas Burger and Jean-Christophe Pesquet for critical review of the manuscript.

## Author contributions

Paul-Henri Pinart (Conceptualization [lead], Investigation [lead], Methodology [lead], Software [lead], Validation [lead], Writing - Original Draft Preparation [equal], Writing - review editing [equal]), Annelaure Damont (Data curation [lead], Methodology [supporting], Validation [supporting]), Sylvain Dechaumet (Conceptualization [supporting], Software [supporting], Supervision [supporting], Validation [supporting]), and Etienne Thèvenot (Conceptualization [supporting], Project Administration [lead], Supervision [lead], Validation [supporting], Writing - Original Draft Preparation [equal], Writing - review editing [equal]).

## Supplementary data

Supplementary data are available in the Supplementary File. Conflict of interest: None declared.

## Data availability

The code and documentation of the MassGAT model, including the generation of the synthetic data, the annotation of the pseudo-spectra, and the application to new data, are publicly available at https://github.com/odisce/MassGAT.

## References

S. Alseekh, A. Aharoni, Y. Brotman, K. Contrepois, J. D’Auria, J. Ewald, J. C. Ewald, P. D. Fraser, P. Giavalisco, R. D. Hall, M. Heinemann, H. Link, J. Luo, S. Neumann, J. Nielsen, L. Perez de Souza, K. Saito, U. Sauer, F. C. Schroeder, S. Schuster, G. Siuzdak, A. Skirycz, L. W. Sumner, M. P. Snyder, H. Tang, T. Tohge, Y. Wang, W. Wen, S. Wu, G. Xu, N. Zamboni, and A. R. Fernie. Mass spectrometry-based metabolomics: A guide for annotation, quantification and best reporting practices. Nature Methods, 18(7):747–756, 2021 July. ISSN 1548-7105. doi: 10.1038/s41592-021-01197-1.

S. Böcker, M. C. Letzel, Z. Liptak, and A. Pervukhin. SIRIUS: Decomposing isotope patterns for metabolite identification. Bioinformatics, 25(2):218–224, 2009. doi: 10.1093/bioinformatics/btn603.

S. Boudah, M.-F. Olivier, S. Aros-Calt, L. Oliveira, F. Fenaille, J.-C. Tabet, and C. Junot. Annotation of the human serum metabolome by coupling three liquid chromatography methods to high-resolution mass spectrometry. Journal of Chromatography B, 966:34–47, Sept. 2014. ISSN 1570-0232. doi: 10.1016/j.jchromb.2014.04.025.

X. Bresson and T. Laurent. Residual gated graph convnets. arXiv preprint arXiv:1711.07553, 2017.

C. Bueschl, M. Doppler, E. Varga, B. Seidl, M. Flasch, B. Warth, and J. Zanghellini. PeakBot: Machine-learning-based chromatographic peak picking. Bioinformatics, 38 (13):3422–3428, 2022 July. ISSN 1367-4803. doi: 10.1093/bioinformatics/btac344.

F. A. Castelli, G. Rosati, C. Moguet, C. Fuentes, J. Marrugo-Ramĺrez, T. Lefebvre, H. Volland, A. MerkoÇi, S. Simon, F. Fenaille, and C. Junot. Metabolomics for personalized medicine: The input of analytical chemistry from biomarker discovery to point-of-care tests. Analytical and Bioanalytical Chemistry, 414:759–789, 2021. ISSN 1618-2642, 1618-2650. doi: 10.1007/s00216-021-03586-z.

L. Chen, W. Lu, L. Wang, X. Xing, Z. Chen, X. Teng, X. Zeng, D. Muscarella, Y. Shen, A. Cowan, M. R. McReynolds, B. J. Kennedy, A. M. Lato, S. R. Campagna, M. Singh, and J. D. Rabinowitz. Metabolite discovery through global annotation of untargeted metabolomics data. Nature Methods, 18(11):1377–1385, 2021. ISSN 1548-7105. doi: 10.1038/s41592-021-01303-3.

B. C. DeFelice, S. S. Mehta, S. Samra, T. Cajka, B. Wancewicz, J. F. Fahrmann, and O. Fiehn. Mass spectral feature list optimizer (MS-FLO): A tool to minimize false positive peak reports in untargeted liquid chromatography-mass spectroscopy (LC-MS) data processing. Analytical Chemistry, 89(6):3250–3255, 2017. ISSN 0003-2700. doi: 10.1021/acs.analchem.6b04372.

A. Delabrière, C. Gianfrotta, S. Dechaumet, A. Damont, T. Hautbergue, P. Roger, E. L. Jamin, O. Puel, C. Junot, F. Fenaille, and E. A. Thèvenot. mineMS2: Annotation of spectral libraries with exact fragmentation patterns. Journal of Cheminformatics, 17(1):111, 2025 July. ISSN 1758-2946. doi: 10.1186/s13321-025-01051-y.

Y. El Abiead, I. Mohanty, S. Xing, A. Rutz, V. Charron-Lamoureux, T. Damiani, W. Lu, G. J. Patti, N. Zamboni, O. Yanes, and P. C. Dorrestein. A Perspective on Unintentional Fragments and Their Impact on the Dark Metabolome, Untargeted Profiling, Molecular Networking, Public Data, and Repository Scale Analysis. JACS Au, 5(12):5828–5850, Dec. 2025. doi: 10.1021/jacsau.5c01063.

Y. Gloaguen, J. A. Kirwan, and D. Beule. Deep Learning-Assisted Peak Curation for Large-Scale LC-MS Metabolomics. Analytical Chemistry, 94(12):4930–4937, Mar. 2022. ISSN 0003-2700. doi: 10.1021/acs.analchem.1c02220.

E. Grushka. Characterization of exponentially modified gaussian peaks in chromatography. Analytical chemistry, 44(11):1733–1738, 1972.

J. Guo and T. Huan. Mechanistic Understanding of the Discrepancies between Common Peak Picking Algorithms in Liquid Chromatography–Mass Spectrometry-Based Metabolomics. Analytical Chemistry, Mar. 2023. ISSN 0003-2700. doi: 10.1021/acs.analchem.2c04887.

J. Guo, S. Shen, S. Xing, Y. Chen, F. Chen, E. M. Porter, H. Yu, and T. Huan. EVA: Evaluation of Metabolic Feature Fidelity Using a Deep Learning Model Trained With Over 25000 Extracted Ion Chromatograms. Analytical Chemistry, 93(36):12181–12186, Sept. 2021. ISSN 0003-2700. doi: 10.1021/acs.analchem.1c01309.

J. Guo, S. Shen, and T. Huan. Paramounter: Direct Measurement of Universal Parameters To Process Metabolomics Data in a “White Box”.Analytical Chemistry, 94(10):4260–4268, Mar. 2022a. ISSN 0003-2700. doi: 10.1021/acs.analchem.1c04758.

J. Guo, H. Yu, S. Xing, and T. Huan. Addressing big data challenges in mass spectrometry-based metabolomics. Chemical Communications, 2022b. ISSN 1364-548X. doi: 10.1039/D2CC03598G.

C. Jaeger, M. Mèret, C. A. Schmitt, and J. Lisec. Compound annotation in liquid chromatography/high-resolution mass spectrometry based metabolomics: Robust adduct ion determination as a prerequisite to structure prediction in electrospray ionization mass spectra. Rapid Communications in Mass Spectrometry, 31(15):1261–1266, 2017. ISSN 1097-0231. doi: 10.1002/rcm.7905.

C. Junot, F. Fenaille, B. Colsch, and F. Becher. High resolution mass spectrometry based techniques at the crossroads of metabolic pathways. Mass Spectrometry Reviews, 33(6): 471–500, 2014. ISSN 1098-2787. doi: 10.1002/mas.21401.

M. Kachman, H. Habra, W. Duren, J. Wigginton, P. Sajjakulnukit, G. Michailidis, C. Burant, and A. Karnovsky. Deep annotation of untargeted LC-MS metabolomics data with Binner. Bioinformatics, 36(6):1801–1806, 2020. ISSN 1367-4803. doi: 10.1093/bioinformatics/btz798.

E. D. Kantz, S. Tiwari, J. D. Watrous, S. Cheng, and M. Jain. Deep neural networks for classification of LC-MS spectral peaks. Analytical Chemistry, 91(19):12407–12413, Oct. 2019. ISSN 0003-2700, 1520-6882. doi: 10.1021/acs.analchem.9b02983.

M. Katajamaa and M. Orešič. Data processing for mass spectrometry-based metabolomics. Journal of Chromatography A, 1158(1):318–328, 2007 July. ISSN 0021-9673. doi: 10.1016/j.chroma.2007.04.021.

E. Kenar, H. Franken, S. Forcisi, K. Wörmann, H.-U. Häring, R. Lehmann, P. Schmitt-Kopplin, A. Zell, and O. Kohlbacher. Automated Label-free Quantification of Metabolites from Liquid Chromatography–Mass Spectrometry Data. Molecular & Cellular Proteomics, 13(1):348–359, Jan. 2014. ISSN 1535-9476, 1535-9484. doi: 10.1074/mcp.M113.031278.

C. Kuhl, R. Tautenhahn, C. Böttcher, T. R. Larson, and S. Neumann. CAMERA: An Integrated Strategy for Compound Spectra Extraction and Annotation of Liquid Chromatography/Mass Spectrometry Data Sets. Analytical Chemistry, 84(1):283–289, Jan. 2012. ISSN 0003-2700. doi: 10.1021/ac202450g.

R. La Rocca, A. Cioppa, E. Ferrarini, M. Höfte, M. Van Droogenbroeck, E. De Pauw, G. Eppe, and L. Quinton. Relational Graph Convolutional Network for Robust Mass Spectrum Classification. Journal of the American Society for Mass Spectrometry, 36(10):2036–2047, Oct. 2025. ISSN 1044-0305. doi: 10.1021/jasms.5c00055.

S. Li and S. Zheng. Generalized Tree Structure to Annotate Untargeted Metabolomics and Stable Isotope Tracing Data. Analytical Chemistry, 95(15):6212–6217, Apr. 2023. ISSN 0003-2700. doi: 10.1021/acs.analchem.2c05810.

S. Li, A. Siddiqa, M. Thapa, Y. Chi, and S. Zheng. Trackable and scalable LC-MS metabolomics data processing using asari. Nature Communications, 14(1):4113, 2023 July. ISSN 2041-1723. doi: 10.1038/s41467-023-39889-1.

Z. Li, Y. Lu, Y. Guo, H. Cao, Q. Wang, and W. Shui. Comprehensive evaluation of untargeted metabolomics data processing software in feature detection, quantification and discriminating marker selection. Analytica Chimica Acta, 1029:50–57, 2018. ISSN 0003-2670. doi: 10.1016/j.aca.2018.05.001.

N. G. Mahieu, J. L. Spalding, S. J. Gelman, and G. J. Patti. Defining and Detecting Complex Peak Relationships in Mass Spectral Data: The Mz.unity Algorithm. Analytical Chemistry, 88(18):9037–9046, 2016. ISSN 0003-2700. doi: 10.1021/acs.analchem.6b01702.

A. D. Melnikov, Y. P. Tsentalovich, and V. V. Yanshole. Deep learning for the precise peak detection in high-resolution LC-MS data. Analytical Chemistry, 92:588–592, Dec. 2019. ISSN 0003-2700, 1520-6882. doi: 10.1021/acs.analchem.9b04811.

O. D. Myers, S. J. Sumner, S. Li, S. Barnes, and X. Du. Detailed Investigation and Comparison of the XCMS and MZmine 2 Chromatogram Construction and Chromatographic Peak Detection Methods for Preprocessing Mass Spectrometry Metabolomics Data. Analytical Chemistry, 89(17):8689–8695, 2017a. ISSN 0003-2700. doi: 10.1021/acs.analchem.7b01069.

O. D. Myers, S. J. Sumner, S. Li, S. Barnes, and X. Du. One step forward for reducing false positive and false negative compound identifications from mass spectrometry metabolomics data: New algorithms for constructing extracted ion chromatograms and detecting chromatographic peaks. Analytical Chemistry, 89(17):8696–8703, 2017b. ISSN 0003-2700. doi: 10.1021/acs.analchem.7b00947.

W. J. Nash, J. B. Ngere, L. Najdekr, and W. B. Dunn. Characterization of Electrospray Ionization Complexity in Untargeted Metabolomic Studies. Analytical Chemistry, 96 (27):10935–10942, 2024 July. ISSN 0003-2700. doi: 10.1021/acs.analchem.4c00966.

L. M. Petrick and N. Shomron. AI/ML-driven advances in untargeted metabolomics and exposomics for biomedical applications. Cell Reports Physical Science, 3(7):100978, 2022 July. ISSN 2666-3864. doi: 10.1016/j.xcrp.2022.100978.

T. Pluskal, S. Castillo, A. Villar-Briones, and M. Orešič. MZmine 2: Modular framework for processing, visualizing, and analyzing mass spectrometry-based molecular profile data. BMC Bioinformatics, 11(1):395, 2010. ISSN 1471-2105. doi: 10.1186/1471-2105-11-395.

F. Rasche, A. Svatoš, R. K. Maddula, C. Böttcher, and S. Böcker. Computing Fragmentation Trees from Tandem Mass Spectrometry Data. Analytical Chemistry, 83(4):1243–1251, 2011. doi: 10.1021/ac101825k.

G. Renner and M. Reuschenbach. Critical review on data processing algorithms in non-target screening: Challenges and opportunities to improve result comparability. Analytical and Bioanalytical Chemistry, 2023 June. ISSN 1618-2650. doi: 10.1007/s00216-023-04776-7.

A. Roux, Y. Xu, J.-F. Heilier, M.-F. Olivier, E. Ezan, J.-C. Tabet, and C. Junot. Annotation of the Human Adult Urinary Metabolome and Metabolite Identification Using Ultra High Performance Liquid Chromatography Coupled to a Linear Quadrupole Ion Trap-Orbitrap Mass Spectrometer. Analytical Chemistry, 84(15):6429–6437, Aug. 2012. ISSN 0003-2700. doi: 10.1021/ac300829f.

S. Samino, M. Vinaixa, M. Dĺaz, A. Beltran, M. A. Rodrĺguez, R. Mallol, M. Heras, A. Cabre, L. Garcia, N. Canela, et al. Metabolomics reveals impaired maturation of hdl particles in adolescents with hyperinsulinaemic androgen excess. Scientific Reports, 5(1):11496, 2015.

R. Schmid, D. Petras, L.-F. Nothias, M. Wang, A. T. Aron, A. Jagels, H. Tsugawa, J. Rainer, M. Garcia-Aloy, K. Dührkop, A. Korf, T. Pluskal, Z. Kamenĺk, A. K. Jarmusch, A. M. Caraballo-Rodrĺguez, K. C. Weldon, M. Nothias-Esposito, A. A. Aksenov, A. Bauermeister, A. Albarracin Orio, C. O. Grundmann, F. Vargas, I. Koester, J. M. Gauglitz, E. C. Gentry, Y. Hövelmann, S. A. Kalinina, M. A. Pendergraft, M. Panitchpakdi, R. Tehan, A. Le Gouellec, G. Aleti, H. Mannochio Russo, B. Arndt, F. Hübner, H. Hayen, H. Zhi, M. Raffatellu, K. A. Prather, L. I. Aluwihare, S. Böcker, K. L. McPhail, H.-U. Humpf, U. Karst, and P. C. Dorrestein. Ion identity molecular networking for mass spectrometry-based metabolomics in the GNPS environment. Nature Communications, 12(1):3832, 2021 June. ISSN 2041-1723. doi: 10.1038/s41467-021-23953-9.

O. Senan, A. Aguilar-Mogas, M. Navarro, J. Capellades, L. Noon, D. Burks, O. Yanes, R. Guimerà, and M. Sales-Pardo. CliqueMS: A computational tool for annotating in-source metabolite ions from LC-MS untargeted metabolomics data based on a coelution similarity network. Bioinformatics, 35 (20):4089–4097, Oct. 2019. ISSN 1367-4803, 1460-2059. doi: 10.1093/bioinformatics/btz207.

C. A. Smith, E. J. Want, G. O’Maille, R. Abagyan, and G. Siuzdak. XCMS: Processing mass spectrometry data for metabolite profiling using nonlinear peak alignment, matching, and identification. Analytical Chemistry, 78(3):779–787, Feb. 2006. ISSN 0003-2700. doi: 10.1021/ac051437y.

T. Stricker, R. Bonner, F. Lisacek, and G. Hopfgartner. Adduct annotation in liquid chromatography/high-resolution mass spectrometry to enhance compound identification. Analytical and Bioanalytical Chemistry, 413(2):503–517, Jan. 2021. ISSN 1618-2650. doi: 10.1007/s00216-020-03019-3.

T. Suvitaival, S. Rogers, and S. Kaski. Stronger findings from mass spectral data through multi-peak modeling. BMC Bioinformatics, 15(1):208, 2014. ISSN 1471-2105. doi: 10.1186/1471-2105-15-208.

R. Tautenhahn, C. Böttcher, and S. Neumann. Highly sensitive feature detection for high resolution LC/MS. BMC Bioinformatics, 9(1):504, Nov. 2008. ISSN 1471-2105. doi: 10.1186/1471-2105-9-504.

H. Tsugawa, T. Cajka, T. Kind, Y. Ma, B. Higgins, K. Ikeda, M. Kanazawa, J. VanderGheynst, O. Fiehn, and M. Arita. MS-DIAL: Data-independent MS/MS deconvolution for comprehensive metabolome analysis. Nature Methods, 12 (6):523–526, 2015. ISSN 1548-7091. doi: 10.1038/nmeth.3393.

K. Uppal, D. I. Walker, and D. P. Jones. xMSannotator: An R Package for Network-Based Annotation of High-Resolution Metabolomics Data. Analytical Chemistry, 89(2):1063–1067, Jan. 2017. ISSN 0003-2700. doi: 10.1021/acs.analchem.6b01214.

P. Veličković, G. Cucurull, A. Casanova, A. Romero, P. Lio, and Y. Bengio. Graph attention networks. arXiv preprint arXiv:1710.10903, 2017.

J. Wandy, R. Daly, R. Breitling, and S. Rogers. Incorporating peak grouping information for alignment of multiple liquid chromatography-mass spectrometry datasets. Bioinformatics, 31(12):1999–2006, 2015 June. ISSN 1367-4803. doi: 10.1093/bioinformatics/btv072.

R. Wang, M. Lu, S. An, J. Wang, and C. Yu. 3D-MSNet: A point cloud based deep learning model for untargeted feature detection and quantification in profile LC-HRMS data. Bioinformatics, 39(5):btad195, 2023. ISSN 1367-4811. doi: 10.1093/bioinformatics/btad195.

H. Weisser, S. Nahnsen, J. Grossmann, L. Nilse, A. Quandt, H. Brauer, M. Sturm, E. Kenar, O. Kohlbacher, R. Aebersold, and L. Malmström. An Automated Pipeline for High-Throughput Label-Free Quantitative Proteomics. Journal of Proteome Research, 12(4):1628–1644, Apr. 2013. ISSN 1535-3893. doi: 10.1021/pr300992u.

D. S. Wishart. Metabolomics for investigating physiological and pathophysiological processes. Physiological Reviews, 99(4): 1819–1875, 2019. ISSN 0031-9333, 1522-1210. doi: 10.1152/physrev.00035.2018.

M. Yu and V. Philip. Mzrtsim: Raw Data Simulation for Reproducible Gas/Liquid Chromatography–Mass Spectrometry-Based Nontargeted Metabolomics Data Analysis. Analytical Chemistry, 97(32):17309–17314, Aug. 2025. ISSN 0003-2700. doi: 10.1021/acs.analchem.5c01213.

H. Zhang, Z. Xu, X. Fan, Y. Wang, Q. Yang, J. Sun, M. Wen, X. Kang, Z. Zhang, and H. Lu. Fusion of quality evaluation metrics and convolutional neural network representations for roi filtering in lc–ms. Analytical chemistry, 95(2):612–620, 2023.

Z. Zhang, H. Yang, Y. Wang, L. Zhang, and S.-H. Lin. QuanFormer: A Transformer-Based Precise Peak Detection and Quantification Tool in LC-MS-Based Metabolomics. Analytical Chemistry, 2025. ISSN 0003-2700. doi: 10.1021/acs.analchem.4c04531.

F. Zhao, S. Huang, and X. Zhang. High sensitivity and specificity feature detection in liquid chromatography–mass spectrometry data: A deep learning framework. Talanta, 222:121580, 2021. ISSN 0039-9140. doi: 10.1016/j.talanta.2020.121580.

