## Supplementary for "MassGAT: a graph-based collective learning approach for untargeted detection and annotation of LC-MS data"

#### Contents

**Note S1: Supplementary methods.** 2

**Note S2: Training performances of MassGAT.** 6

**Figure S1: MassGAT filtering size and reproducibility.** 7

**Figure S2: Selected peaks per pseudo-spectrum.** 8

**Figure S3: Comparison of the target compounds detected by XCMS, 3D-MSNet and MassGAT.** 9

**Figure S4: Annotated metabolites from the IRS2-KO sample (positive ionization).** 10

**Figure S5: Annotated metabolites from the IRS2-KO sample (negative ionization).** 18

**Figure S6: Re-annotation of Aspartic acid as N-Acetylaspartic acid.** 24

**Figure S7: MassGAT pseudo-spectra of L-Methionine-oxide in the HIAE cohort.** 25

**Table S1: List of m/z differences used to build the components.** 28

**Table S2: Parameter values used to benchmark the data processing software (TripleTOF 6600 and QE HF datasets).** 30

**Table S3: Annotations from CliqueMS and MassGAT on the IRS2-KO sample (positive ionization).** 31

**Table S4: Annotations from CliqueMS and MassGAT on the IRS2-KO dataset (negative ionization).** 33

**Table S5: Annotations from CliqueMS and MassGAT on the HIAE cohort (MTBLS103 dataset).** 34

### **Note S1:** Supplementary methods.

#### Hierachical building of the graph component

A first set of candidate pairs is obtained by comparing the m/z distance of segments within a defined retention time window to a list of 5 isotopes ($^{13}C$, $^{15}N$, $^{18}O$, $^{34}S$, and $^{37}Cl$) with a fixed tolerance (by default $5 \cdot 10^{-4}$ Da). Only candidate isotopic pairs with a cosine similarity of their normalised intensity profiles exceeding 0.5 are kept. The process forms a tree of isotopic links in which the root node is either an adduct, a fragment or a dimer. A second set of pairs is then computed similarly between either the root nodes of those trees or isolated nodes, with m/z differences corresponding to common adducts or fragments (e.g., for [X+Na]-[X+H], $\Delta \approx 21.9819$ Da). Finally, a third set of pairs is computed to look for dimers (e.g., [2X+H] with [X+H]). The tolerance in m/z for both adducts and dimers is also $5 \cdot 10^{-4}$ Da and the threshold cosine similarity is set by default to 0.8. The default list of chemical rules (Table S1) and threshold values can be modified by the user to best reflect the experimental setup.

#### Architecture of the GAT model

##### Input encoding

The 72 dimensional encoding vector is split between the meta features and the elution profile. To enable the network to capture high-frequency variations in scalar inputs such as $m/z$ and intensities, we project these meta-features using random Fourier features (Tancik et al., 2020) followed by a two-layer Multilayer Perceptron (MLP), resulting in a $d$-dimensional embedding. For connected nodes the encoder is a single-layer Transformer ($d_{\text{model}} = 64$, 2 attention heads) with attention pooling over the 32 time-points, which captures global shape coherence across the full elution profile. For isolated nodes the encoder is a three-layer 1-D CNN ($2 \to 16 \to 32 \to 64$ channels, kernel sizes 5, 3, 3, followed by adaptive max-pooling). The embeddings from the meta-features and the elution profile are concatenated and projected to the hidden dimension $h = 32$ through a linear layer, giving an initial node representation $\mathbf{x}_v^{(0)} \in \mathbb{R}^h$ for each node $v$.

##### Edge attributes

Each edge attribute vector is obtained by concatenating a 4-dimensional learnt embedding that encodes the semantic type of the chemical relationship (isotope types, adducts, fragments, and dimers) to four other continuous features: the cosine similarity of the elution profiles from the two adjacent nodes, the log-ratio of the integrated intensities, the log-ratio of the maximum intensities, and the retention time difference. This results in a 8-dimensional edge attribute $\mathbf{a}_e \in \mathbb{R}^8$.

##### Gated message passing

The initial embeddings are passed through $K = 2$ blocks, each consisting of $L = 2$ graph attention layers (Veličković et al., 2017) with a learned edge-gating mechanism (Bresson and Laurent, 2017), $g_e \in [0, 1]$, computed for every edge $e = (u, v)$:

$$g_e = \sigma(\text{MLP}_{\text{gate}}([\mathbf{x}_u \,\|\, \mathbf{x}_v \,\|\, \mathbf{a}_e])) \tag{1}$$

where $\sigma$ is the sigmoid function and $\text{MLP}_{\text{gate}}$ maps $(2h + 12)$ dimensions to a scalar through a 64-unit hidden layer. The gate prevents aggregation from noisy nodes into a clean component, something the attention mechanism fails to do on its own. Residual connections are applied within and across blocks to prevent over-smoothing. The isolated nodes, which do not receive any message by definition, keep their initial embedding from the encoder.

After the final block, a shared projection layer maps all node representations to an embedding space. Three prediction heads then run in parallel: a node head classifies each ion as a true peak or noise, an edge head assigns a score to each candidate pair to determine whether or not it belongs to the same metabolite, and a graph head classifies each component (true vs. noise). Separate node heads are used for connected and isolated ions, as initial experiments have shown that a shared head can interfere with the classification of isolated ions.

##### Semi-supervised learning and loss formulation

Three different labels are defined at the node level: 1) nodes confirmed as true peaks have a label $y_v = 1$ common to all nodes from the same metabolite, 2) nodes confirmed as noise have the label $y_v = -1$, and 3) nodes in real partially annotated components with an unknown status have the label $y_v = -2$ and are excluded from all loss terms to avoid penalizing the model for predictions of nodes that were not manually validated. This allows the integration of real MS1 spectra (which are only partially annotated manually). Graph-level labels are generated automatically: a connected component is labeled as positive if it contains at least one node with $y_v \geq 0$.

Nodes are then split into connected nodes (which belong components with at least one edge), and isolated nodes (which form single-node components on their own). The node loss is computed separately for these two groups, with isolated positive nodes down-weighted relative to connected ones, since single-node components are less reliable than validated multi-node components. Isolated nodes are excluded from the graph, edge and consistency loss terms.

The total loss combines four terms:

$$\mathcal{L} = \alpha\,\mathcal{L}_{\text{graph}} + \beta\,\mathcal{L}_{\text{node}} + \gamma\,\mathcal{L}_{\text{edge}} + \delta\,\mathcal{L}_{\text{cons}} \tag{2}$$

The node loss $\mathcal{L}_{\text{node}}$, graph loss $\mathcal{L}_{\text{graph}}$, and edge loss $\mathcal{L}_{\text{edge}}$ are all computed as focal losses (Lin et al., 2017), with $\mathcal{L}_{\text{graph}}$ evaluated only on non-singleton components. A fourth consistency loss is used to enforce coherent predictions from the different heads:

$$\mathcal{L}_{\text{cons}} = \frac{1}{|E|} \sum_{(u,v)\in E} \max\Big(0, \hat{p}_e - \min(\hat{p}_u, \hat{p}_v)\Big) + \frac{1}{G} \sum_{g=1}^{G} \Big[\max\Big(0, \hat{p}_g^{\max} - \hat{p}_g - \epsilon\Big) + \max\Big(0, \hat{p}_g - \hat{p}_g^{\max} - \epsilon\Big)\Big] \tag{3}$$

where $\hat{p}_g^{\max} = \max_{v \in g} \hat{p}_v$ and $\epsilon = 0.1$. The first term enforces the requirement that an edge prediction cannot be greater than the probability of its connected nodes, applying the same hinge-loss relaxation of logical implication (i.e., Node A and Node B implies $\text{Edge}_{A,B}$) from Probabilistic Soft Logic (Bach et al., 2017). The second term penalizes graph probabilities when they are further from the maximum node probability within the component than a margin $\epsilon$, adapting as a soft penalty the max-consistency principle used to enforce label-hierarchy coherence (Giunchiglia and Lukasiewicz, 2020).

##### Implementation

The MassGAT model is implemented in PyTorch (Paszke et al., 2019) and PyTorch Geometric (Fey and Lenssen, 2019). Training uses the Adam optimiser (Kingma and Ba, 2014) with a learning rate of $3 \times 10^{-4}$ and a batch size of 32 for 22 epochs on NVIDIA A40 GPU.

#### Annotation heuristic

Let $\mathcal{G} = (\mathcal{V}, \mathcal{E})$ be a validated connected component. A node $v \in \mathcal{V}$ is considered a *candidate main ion* if it has not been identified as an isotope or dimer child — i.e., it does not appear as the heavier node of any isotope or dimer edge in $\mathcal{E}$. For a given candidate

$v^*$, the undirected edges of $\mathcal{G}$ are assigned a chemical direction to enumerate its valid children and propagate annotations. For isotope edges, the heavier node is systematically assigned as the child. For the other edges, strict directionality is enforced: adducts (e.g. Na-H, $[K\text{-}H]^+$) must propagate towards the heavier node, while losses (e.g. $CO_2$) must propagate towards the lighter node. Dimer edges naturally point to the heavier node; however, a fractional mass defect heuristic ($0.45 \leq m/z \pmod 1 \leq 0.85$) is applied to detect potential multicharged ions (e.g., $[M{+}2H]^{2+}$).

Each candidate $v^*$ is then assigned a normalized score:

$$s(v^*) = 1.0\,\frac{n_{\text{children}}}{\max n_{\text{children}}} + 0.5\,\frac{n_{\text{isotopes}}}{\max n_{\text{isotopes}}} + 2.0\,\frac{I_{v^*}}{\max I_v} + 0.5\,\frac{\hat{p}_{v^*}}{\max \hat{p}_v},$$

where $n_{\text{children}}$ is the total number of children, $n_{\text{isotopes}}$ the number of isotope children, and $I_{v^*}$ the linear integrated intensity. Normalizing these metrics relative to the component's maxima prevents too much variance across different datasets. The contribution of $n_{\text{children}}$ and $n_{\text{isotopes}}$ to the score reflects the observation that the true main ion tends to act as a root in the graph, accumulating the largest number of consistent neighbours. Intensity, and to a lesser extent probability, act as a deciding factor in distinguishing between ions that would otherwise have the same type of neighbourhood.

Given $K$ candidates with scores $s_1, \ldots, s_K$, a softmax distribution with temperature $T = 2$ is computed as

$$\pi_k = \frac{\exp((s_k - \max_j s_j)/T)}{\sum_{j=1}^{K} \exp((s_j - \max_j s_j)/T)}, \tag{4}$$

in order to create a probabilistic ranking over annotation hypotheses. The implied neutral mass $M$ is estimated by weighted regression over all non-dimer nodes.

#### Evaluation of peak detection

For both TripleTOF 6600 and QE HF datasets (Li et al., 2018), the 8 centroided .mzML files were processed independently by MassGAT (parameter values in Table S2). The entire pipeline was completed in 137 s for a TOF file (65 s for the segmentation of ion traces by ionshed, 36 s for the building of candidate graph components, 36 s for graph inference, and 1 s for annotation of the validated pseudo-spectra with less than 200 nodes) and in 502 s for a QE HF file (207 s, 206 s, 89 s, and 3 s, respectively) using 5 CPU cores without GPU.

The peaks present in at least 50% of 8 replicates were then aligned between the samples with the XCMS grouping and fillpeaks procedures (Smith et al., 2006). The consensus peaks (i.e., peaks without missing values in the final peak table) were used to determine the recall (proportion of detected peaks from the ground truth), by using the tolerance values from Li et al., 2018: 10 ppm (respectively, 5 ppm) for the TripleTOF 6600 data (respectively, the QE HF data), and 0.3 min for both.

These recall values were compared to those obtained with the XCMS (Tautenhahn et al., 2008), asari (Li et al., 2023), and 3D-MSNet (Wang et al., 2023) software (parameter values in Table S2). Note that, as 3D-MSNet was trained on one of the files from this dataset, we removed the corresponding file from the evaluation.

#### Evaluation of componentization and annotation

Componentization and annotation were assessed on the Retina IRS2-KO and MTBLS103 datasets, previously used by Senan et al., 2019 to benchmark the performances of the CliqueMS software and the alternative approaches CAMERA (Kuhl et al., 2012), xMSannotator (Uppal et al., 2017), and MS-FLO (DeFelice et al., 2017).

CliqueMS was run as described in the original publication (Senan et al., 2019). To ensure a fair comparison, the default list of adducts and multimers from CliqueMS was modified to include the additional m/z differences from MassGAT.

The performances for xMSannotator, MS-FLO, and CAMERA are those reported from the previous benchmark by Senan et al. 2019.

### **Note S2:** Training performances of MassGAT.

The MassGAT model was trained with a learning rate of $5 \times 10^{-5}$, a batch size of 64, and hidden and embedding dimensions of 32 and 64, respectively. The dataset was split into 3 groups: 70% for training, 20% for validation, and 10% for testing. With the early stopping criterion based on validation loss performance, the training process terminated at epoch 44, with a convergence across the four loss components. The evolution and convergence of these loss terms normalized to their initial values after epoch 1 are shown on the figure below. The rapid decay of the consistency loss (green) demonstrates the successful enforcement of hierarchical logic ensuring that graph-level confidence remains supported by the underlying node probabilities.

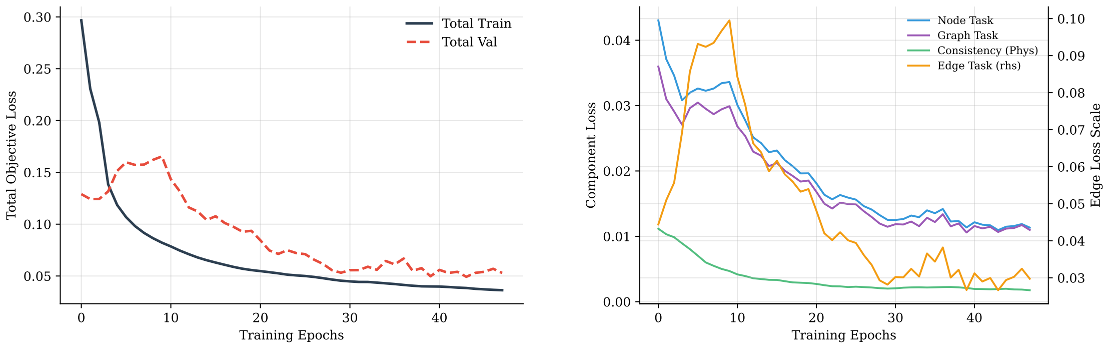


**(a)** **(b)**

The final performances of the model are shown below:

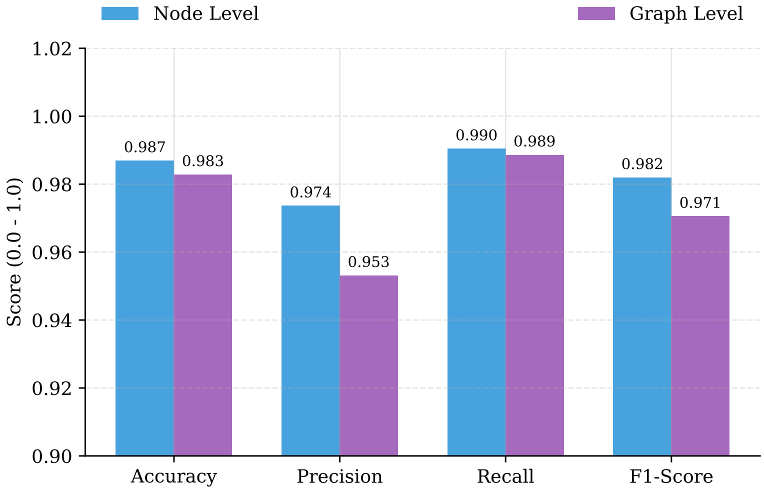

### Figure S1: MassGAT filtering size and reproducibility.

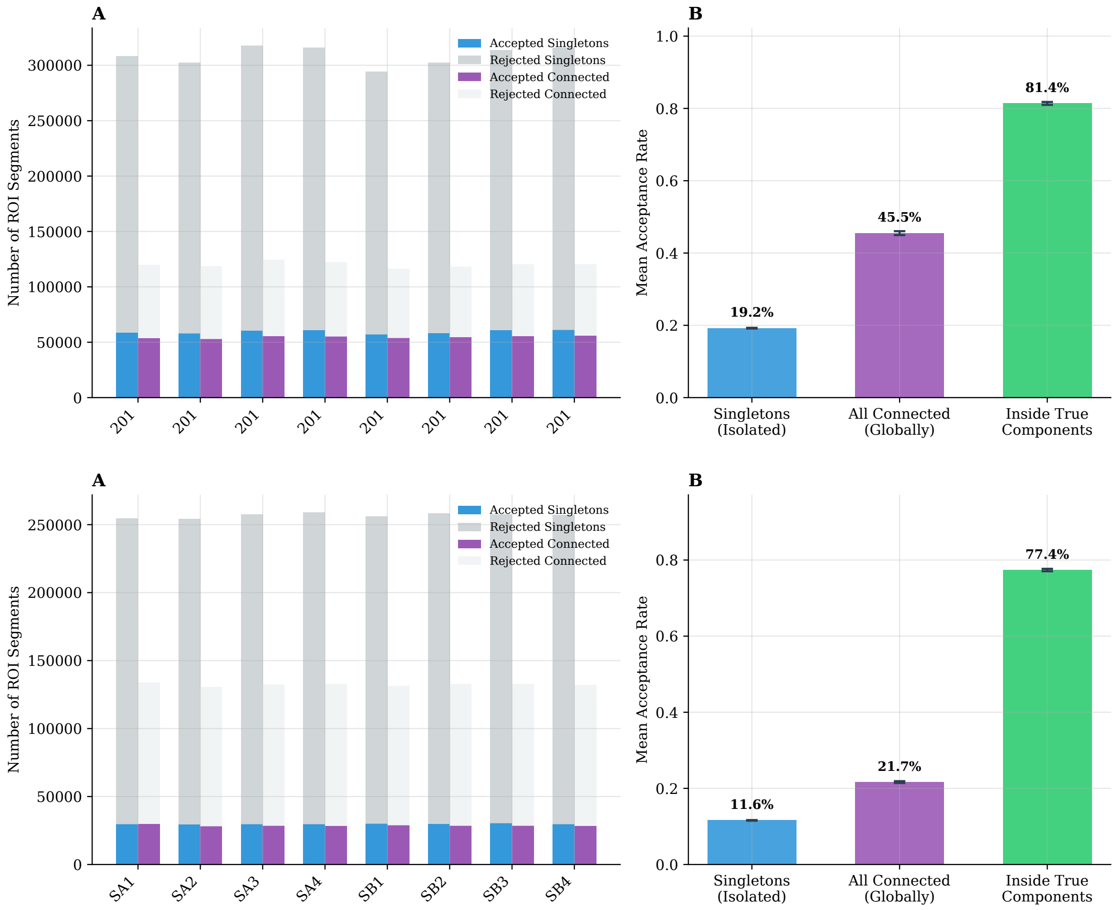


Signal filtering efficiency by MassGAT of the TripleTOF 6600 (top) and QE HF (bottom) samples. **A**: Initial number of segmented ion traces (ROIs) and number of validated peaks for each replicate. **B**: Mean acceptance rates for isolated singletons, all connected traces, and traces specifically located inside true components (with 95% confidence intervals). The data show that connection increases the probability of validation, but pruning still occurs within true components.

### Figure S2: Selected peaks in pseudo-spectra.

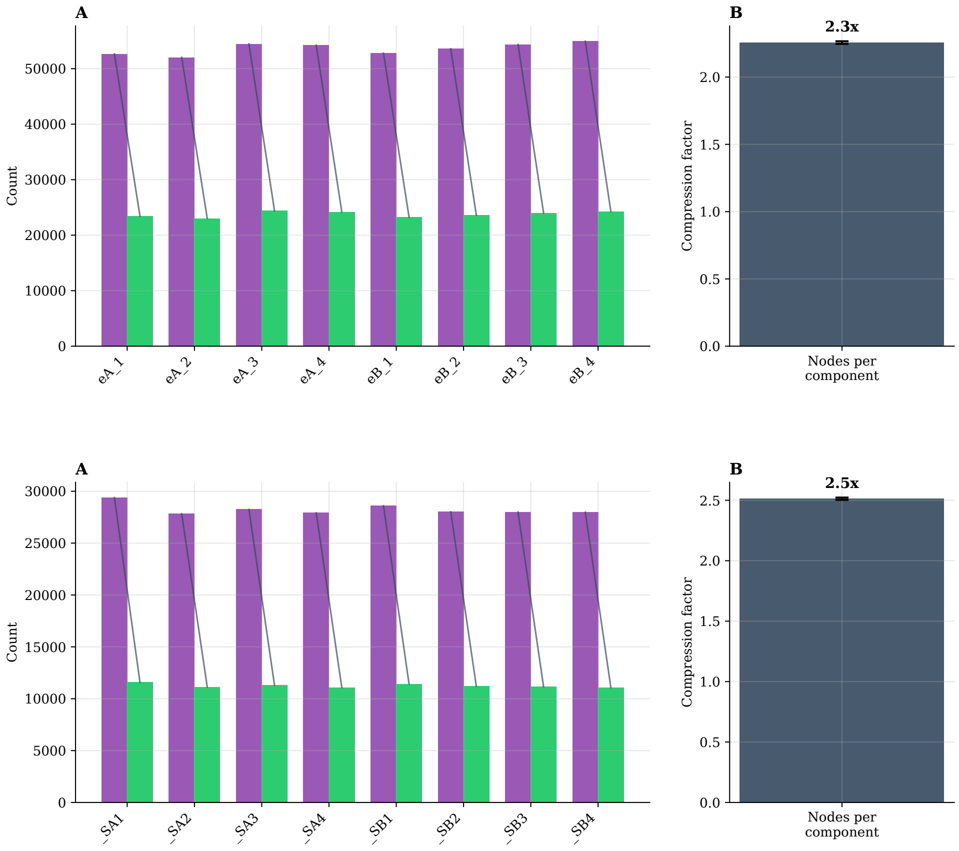


Number of pseudo-spectra and their peaks in the TripleTOF 6600 (top) and QE HF (bottom) samples. **A**: Bars corresponding to pseudo-spectra and peaks are coloured in green and purple, respectively, for each replicate. **B**: Average ratio of peaks per pseudo-spectrum.

**Figure S3:** Comparison of the target compounds detected by XCMS, 3D-MSNet and MassGAT.

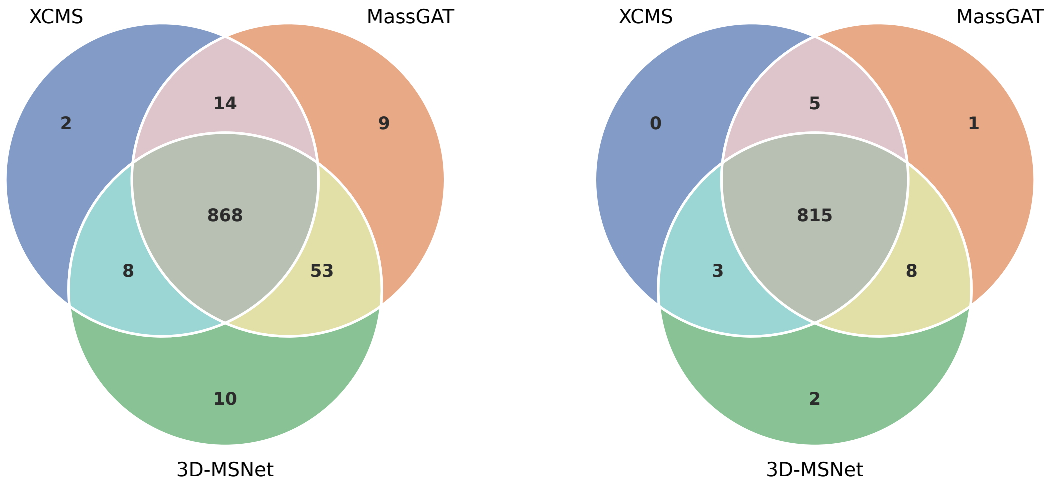


TripleTOF 6600 QE HF

Number of target compounds which are detected by the XCMS, 3D-MSNet and MassGAT software in the benchmark datasets (Li et al., 2018) TripleTOF 6600 (left) and QE HF (right).

### **Figure S4:** Annotated metabolites from the IRS2-KO sample (positive ionization).

The pseudo-spectra from the annotated metabolites (i.e., with at least two nodes) described in Table S3 are visualized below. The star corresponds to the targeted ground-truth peak. Nodes (detected ions) are coloured according to their detection probability (green: 1, yellow: 0.5). Edge are labeled by the chemical formula of the m/z difference, and coloured according to type of link (green: isotope, blue: adduct, purple: loss, brown: loss or neutral adduct, orange: dimer or doubly charged).

#### **Metabolites annotated by MassGAT but not CliqueMS (n = 4)**

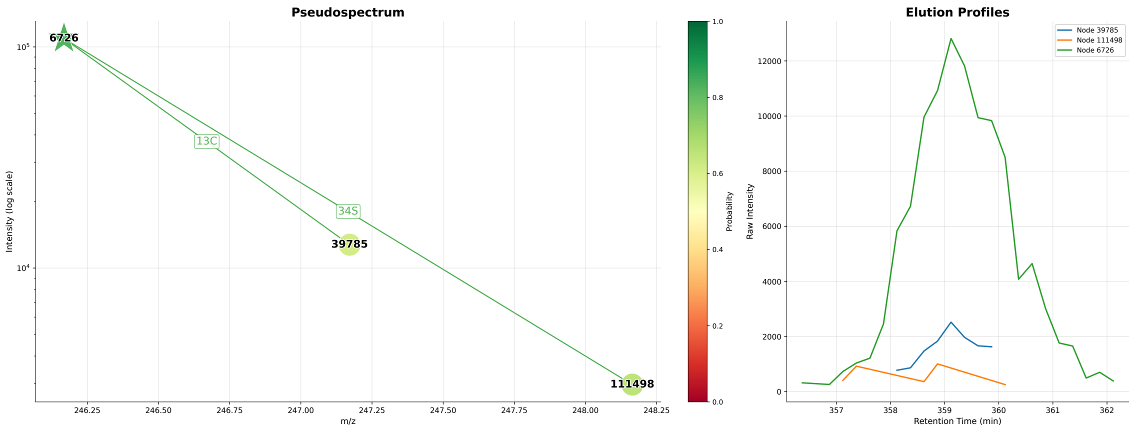


2-Methylbutyroylcarnitine

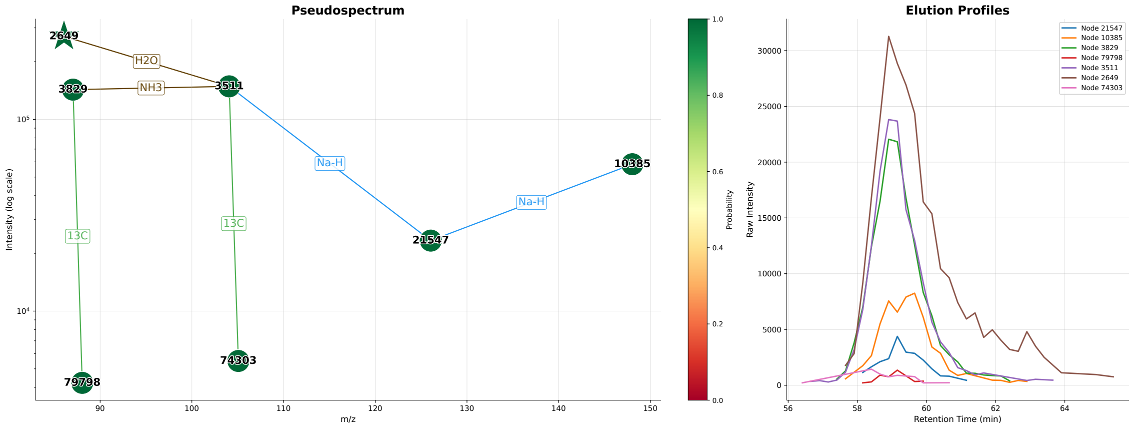


2-Pyrrolidinone

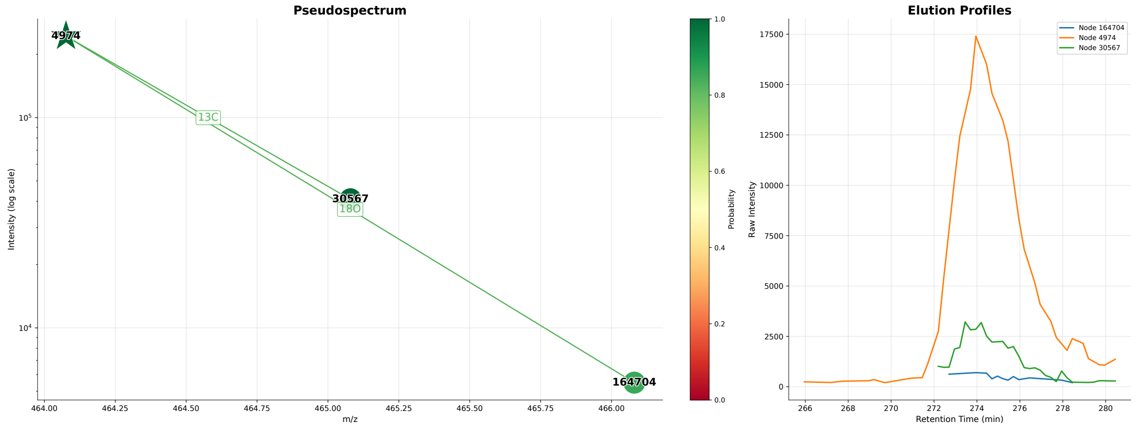


Adenylosuccinic acid

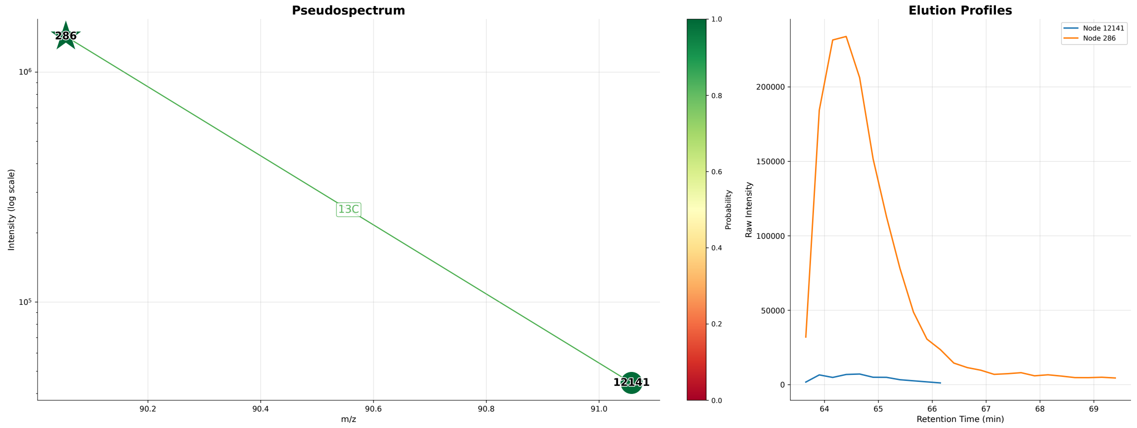


L-Alanine

### Metabolites annotated by MassGAT and CliqueMS (n = 16)

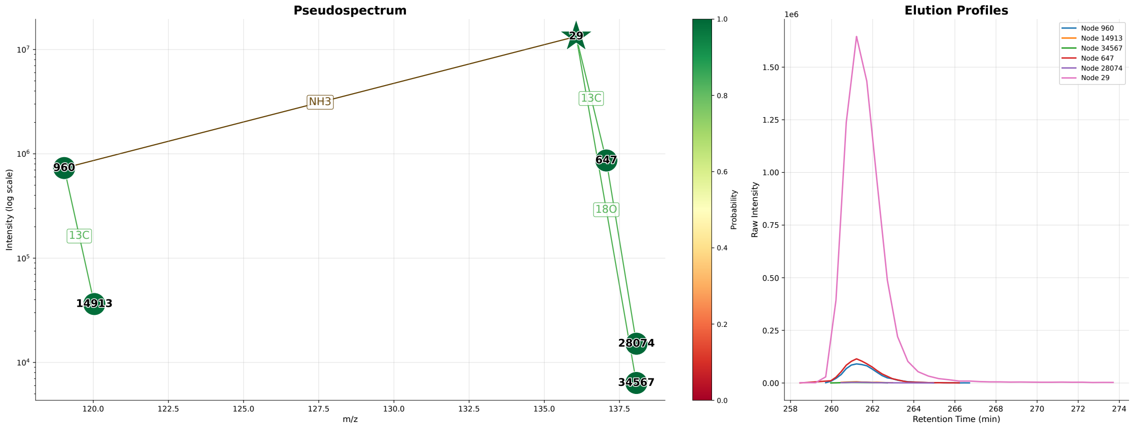


Adenine

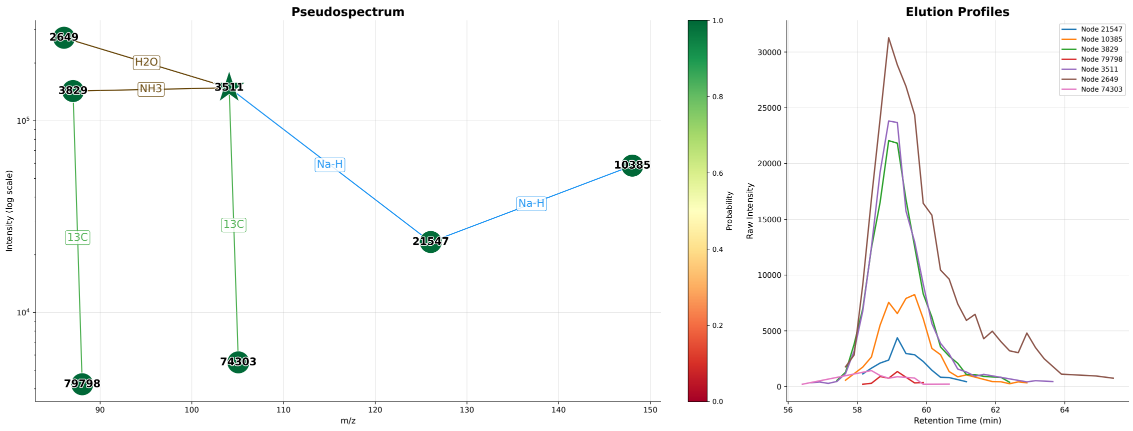


GABA

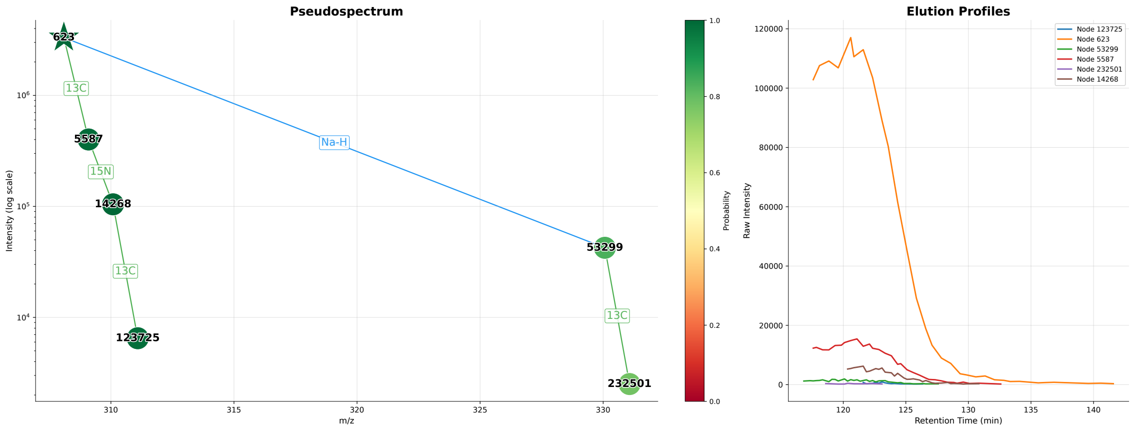


Glutathione

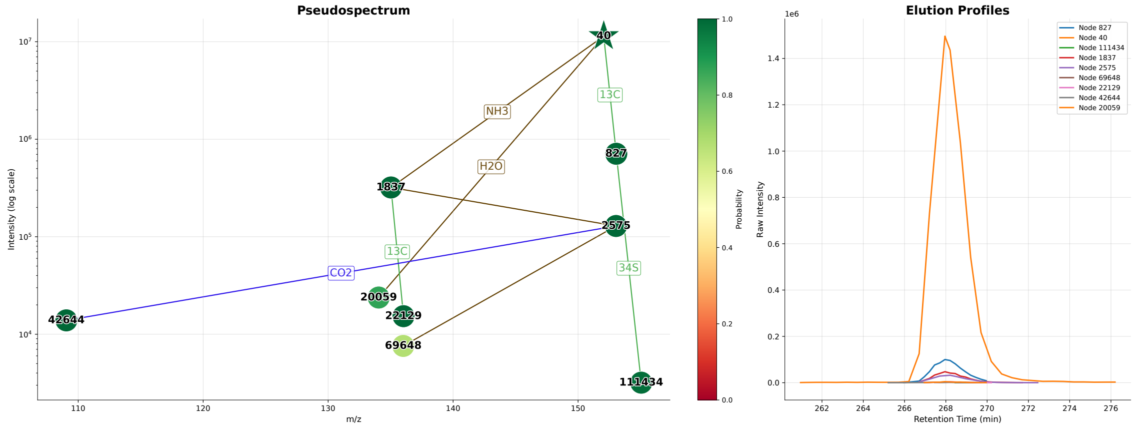


Guanine

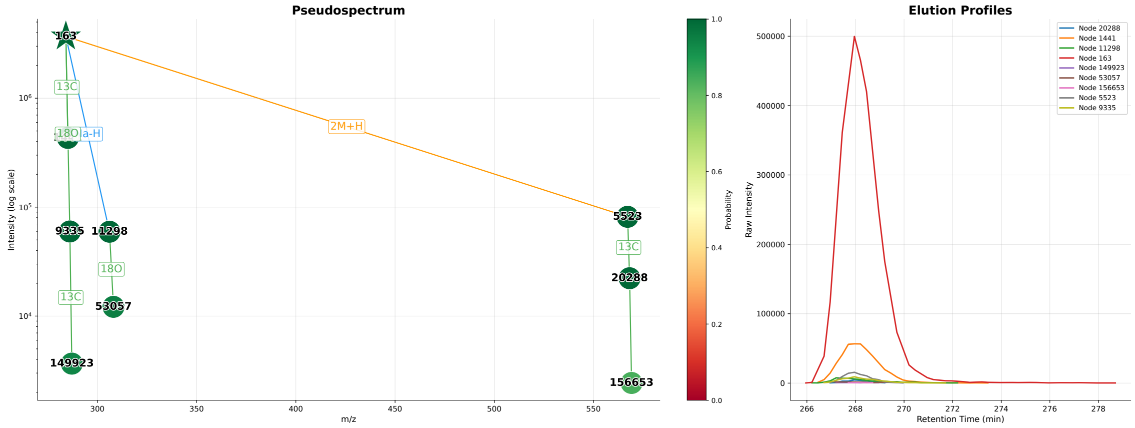


Guanosine

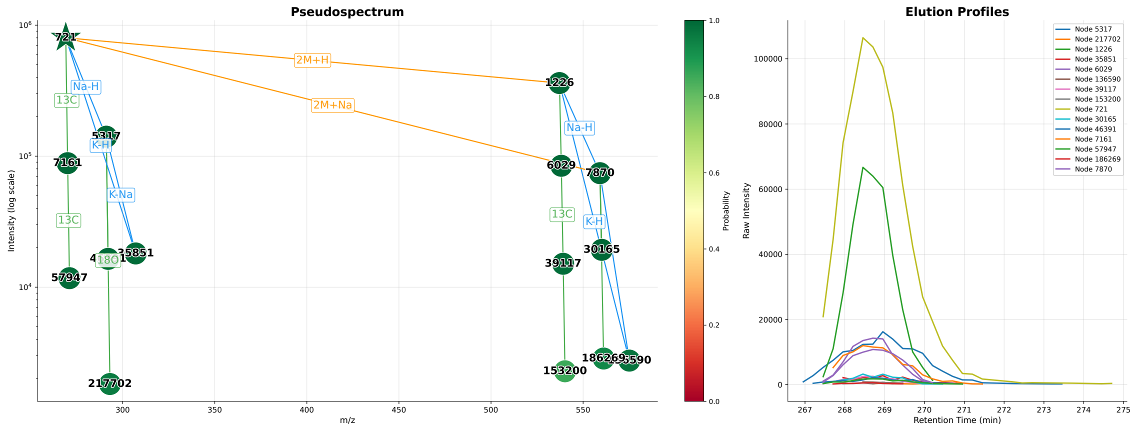


Inosine

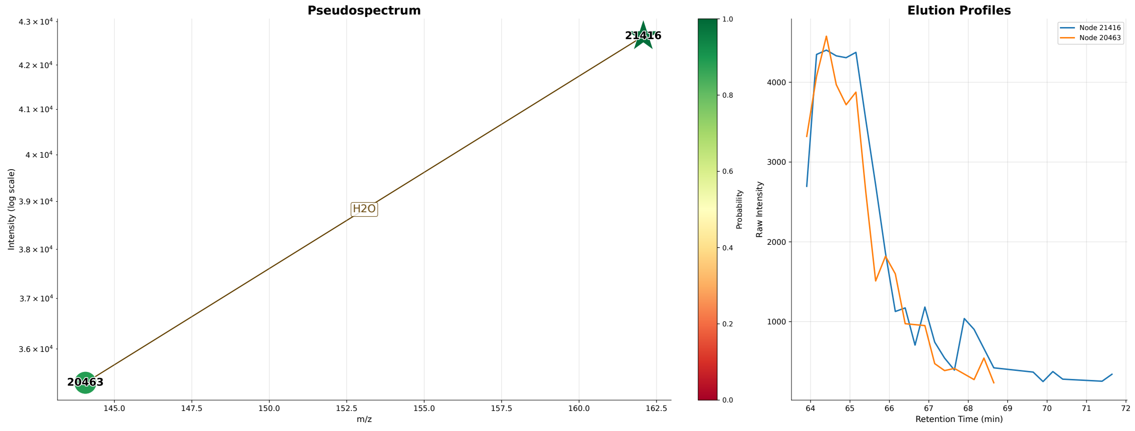


L-2-Aminoadipic acid

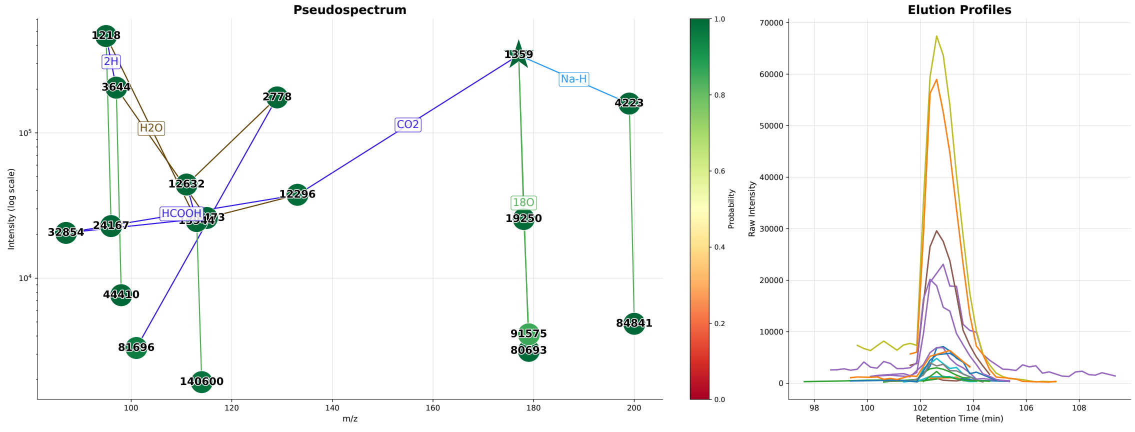


L-Ascorbic acid

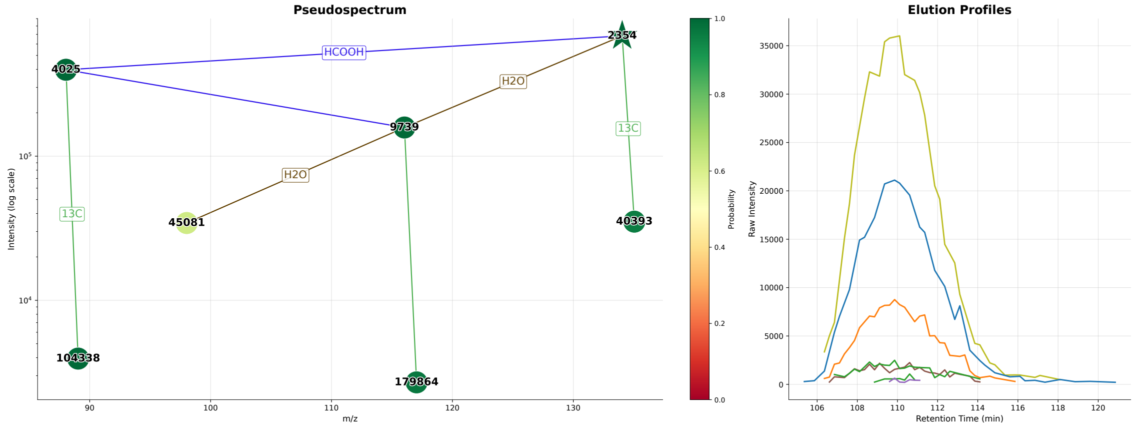


L-Aspartic acid

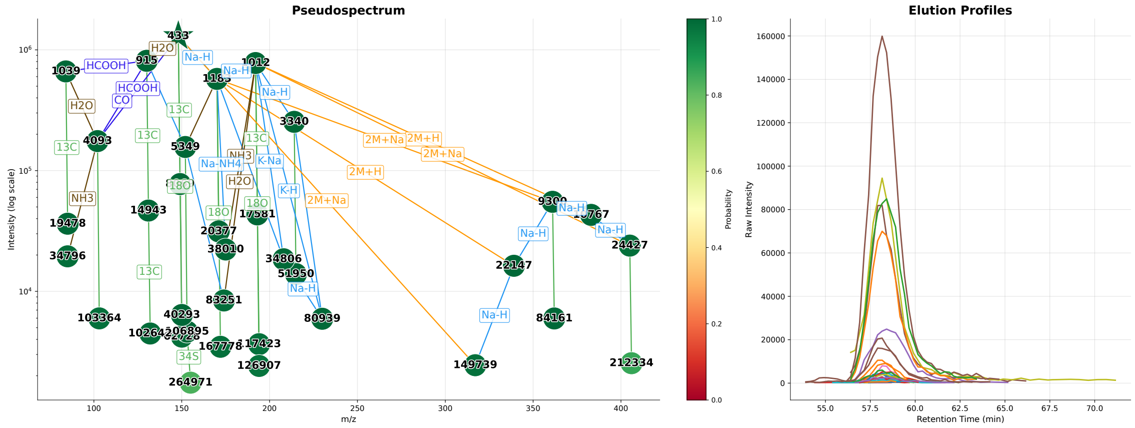


L-Glutamic acid

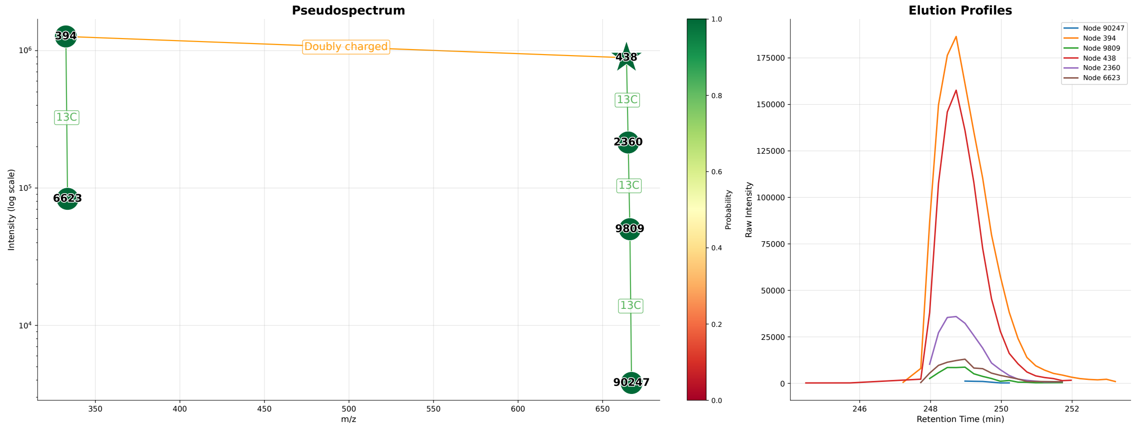


NAD

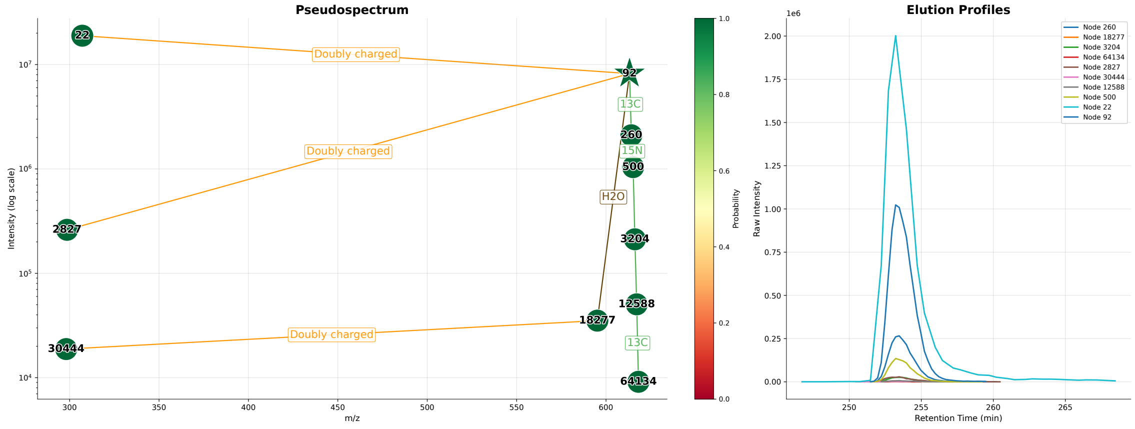


Oxiglutathione

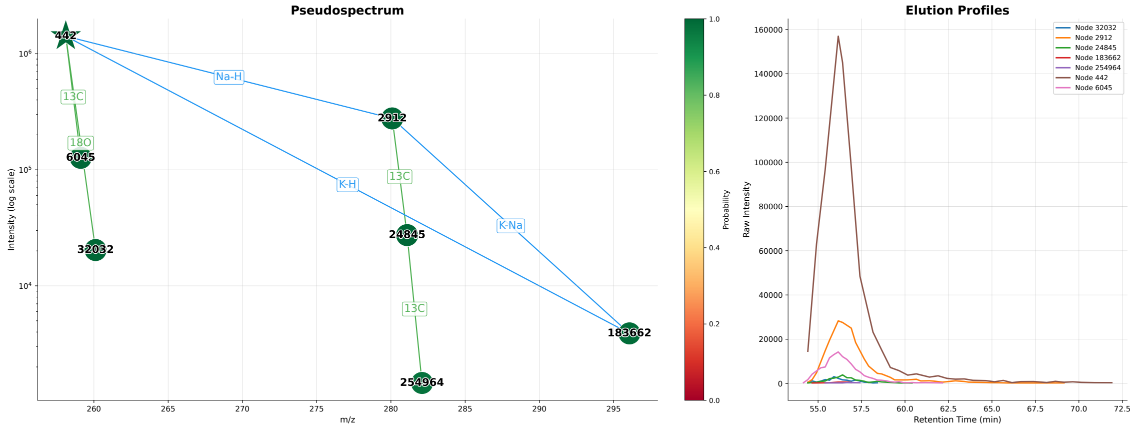


PC

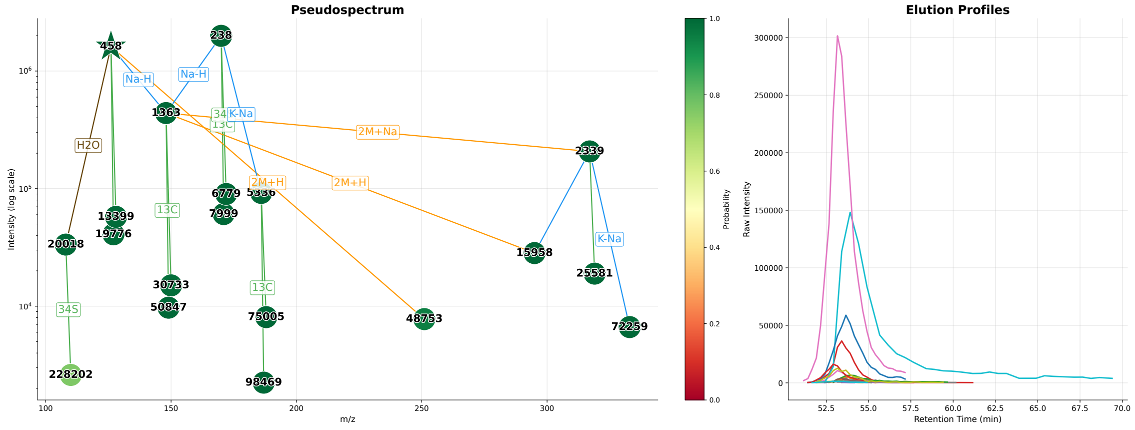


Taurine

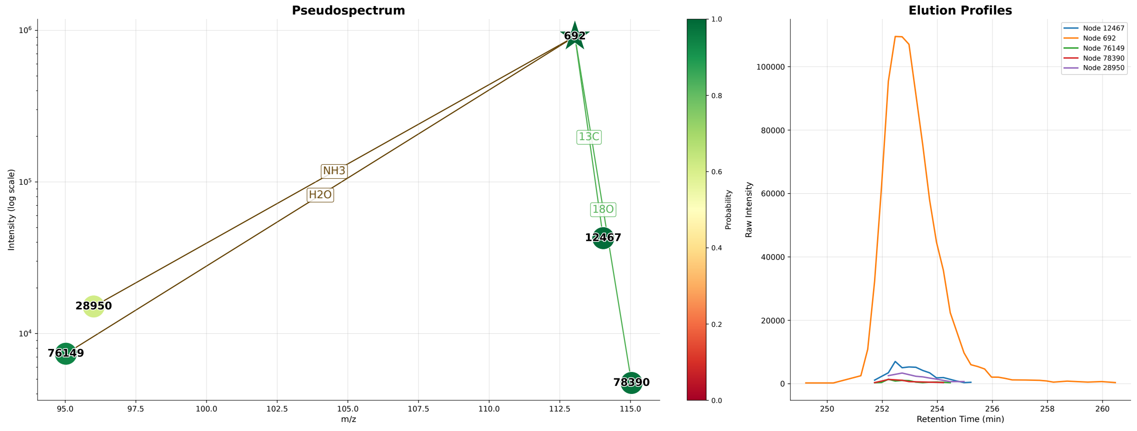


Uracil

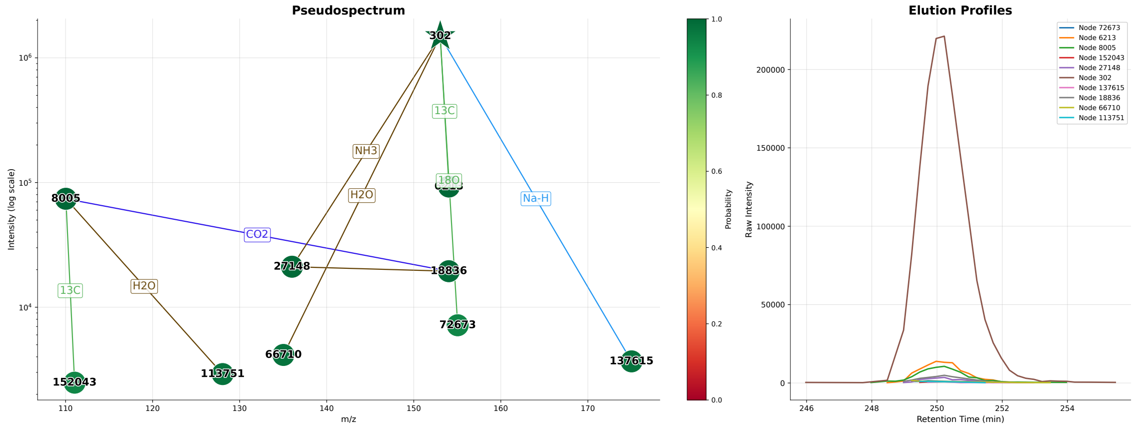
Pseudospectrum
Intensity (log scale)
m/z
302
8005
27148
18836
72673
66710
113751
152043
137615
NH3
H2O
13C
18O
CO2
Na-H
Probability
Elution Profiles
Raw Intensity
Retention Time (min)
Node 72673
Node 6213
Node 8005
Node 152043
Node 27148
Node 302
Node 137615
Node 18836
Node 66710
Node 113751

Xanthine

### **Figure S5:** Annotated metabolites from the IRS2-KO sample (negative ionization).

The pseudo-spectra from the annotated metabolites (i.e., with at least two nodes) described in Table S4 are visualized below. The star corresponds to the targeted ground-truth peak. Nodes (detected ions) are coloured according to their detection probability (green: 1, yellow: 0.5). Edge are labeled by the chemical formula of the m/z difference, and coloured according to type of link (green: isotope, blue: adduct, purple: loss, brown: loss or neutral adduct, orange: dimer or doubly charged).

#### **Metabolites annotated by MassGAT but not CliqueMS (n = 11)**

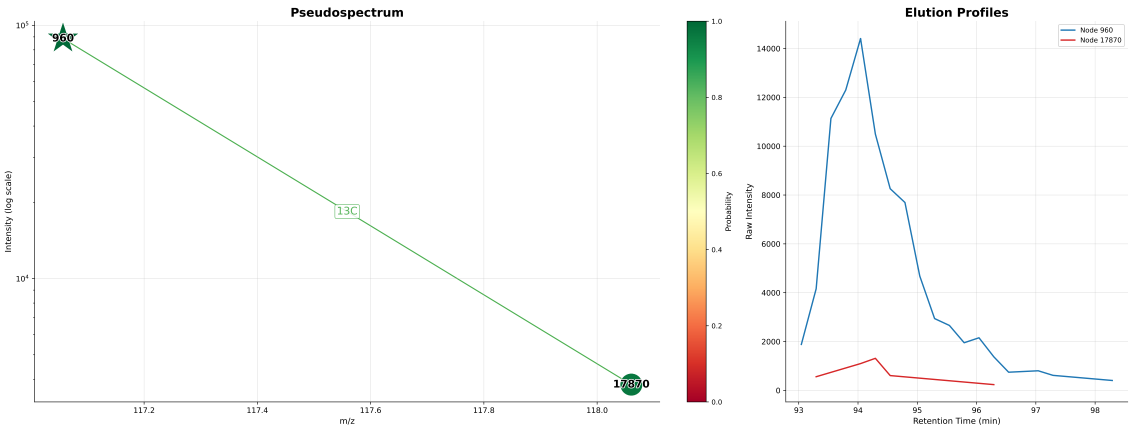


2-hydroxy-2-methyl-butyric acid

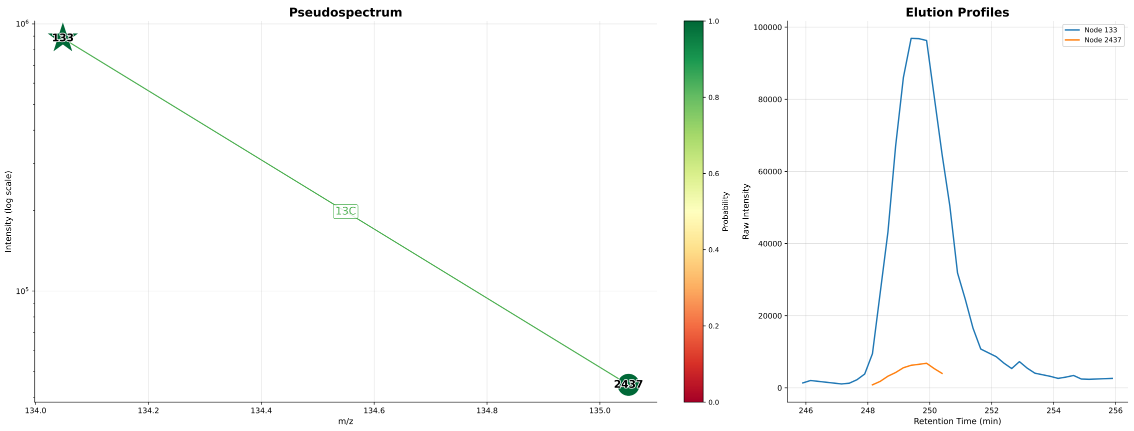


Adenine

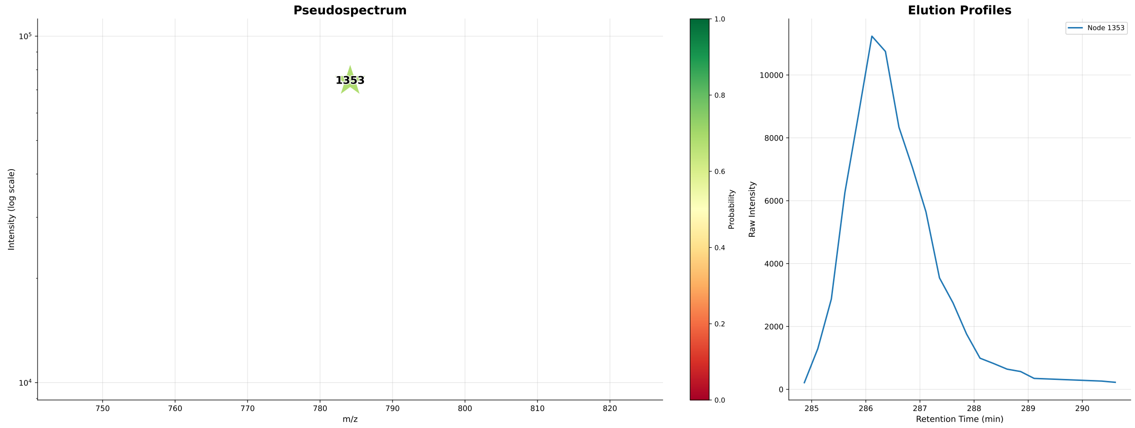


FAD

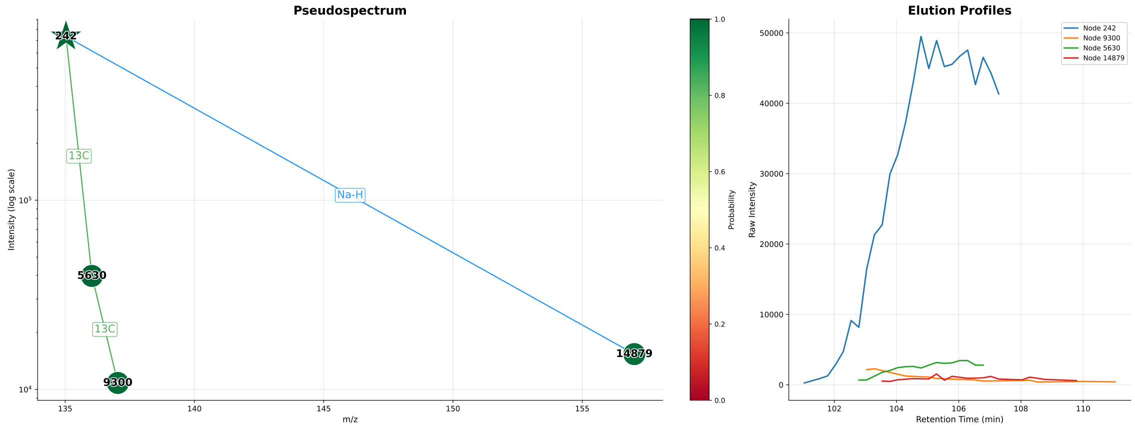


Hypoxanthine

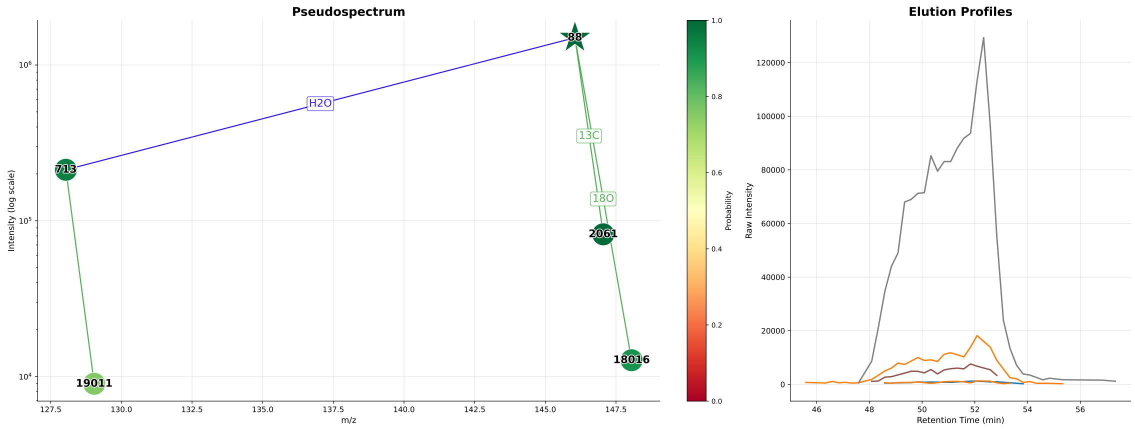


L-Glutamic acid

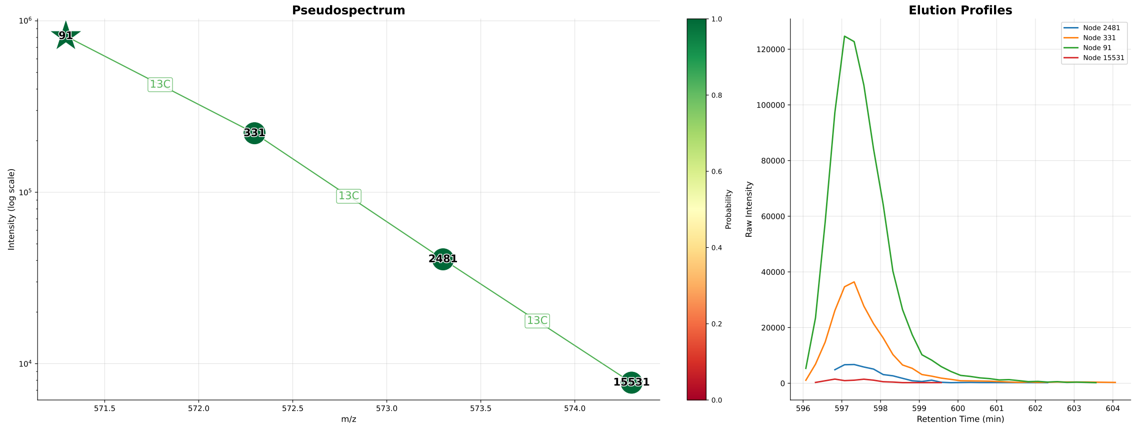


LPI I

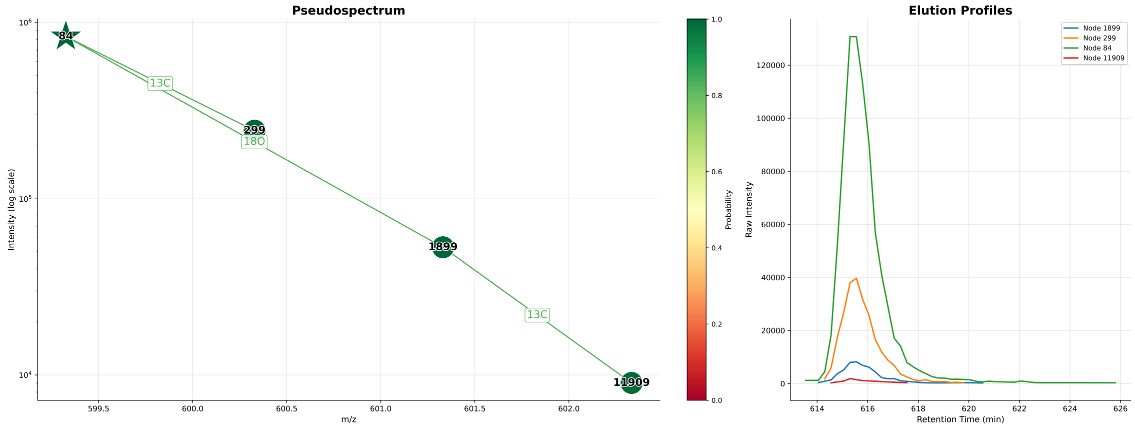


LPI II

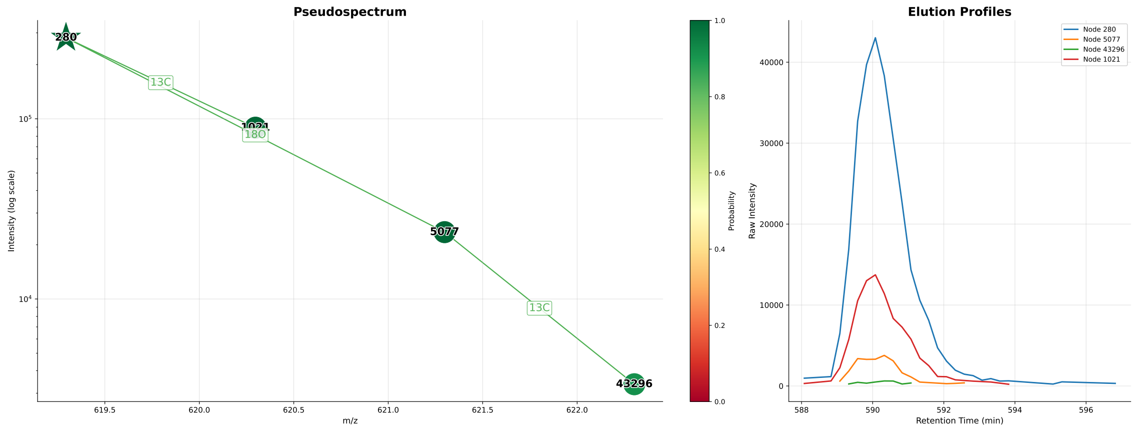


LPI III

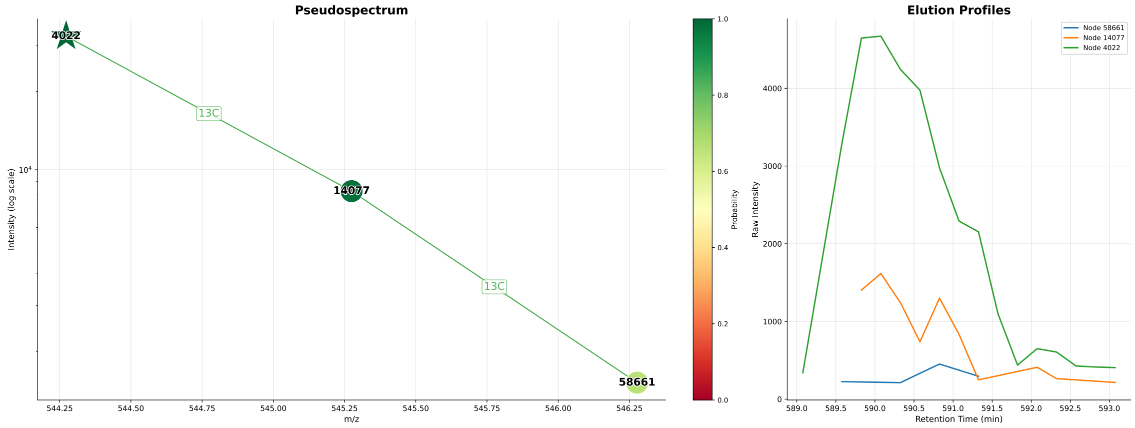


LPSE I

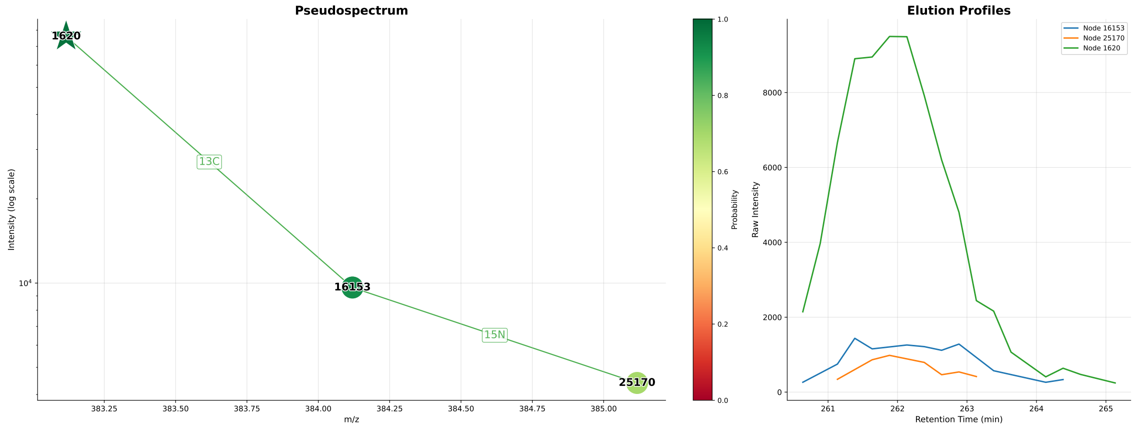


S-Adenosylhomocysteine

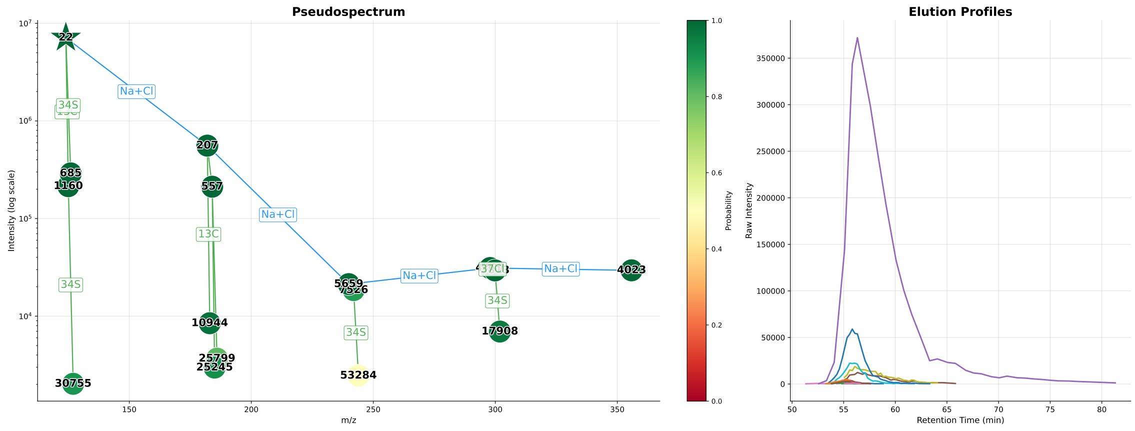


Taurine

### Metabolites annotated by MassGAT and CliqueMS (n = 6)

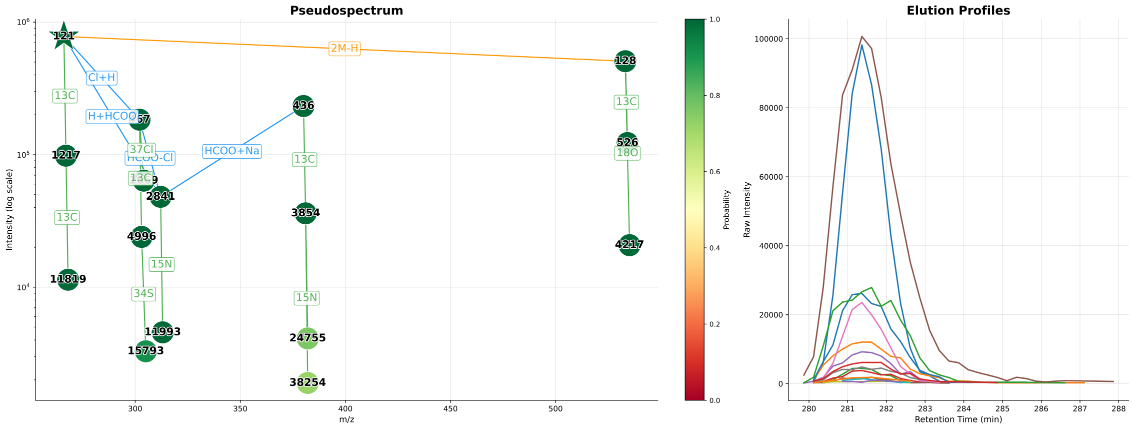


Adenosine

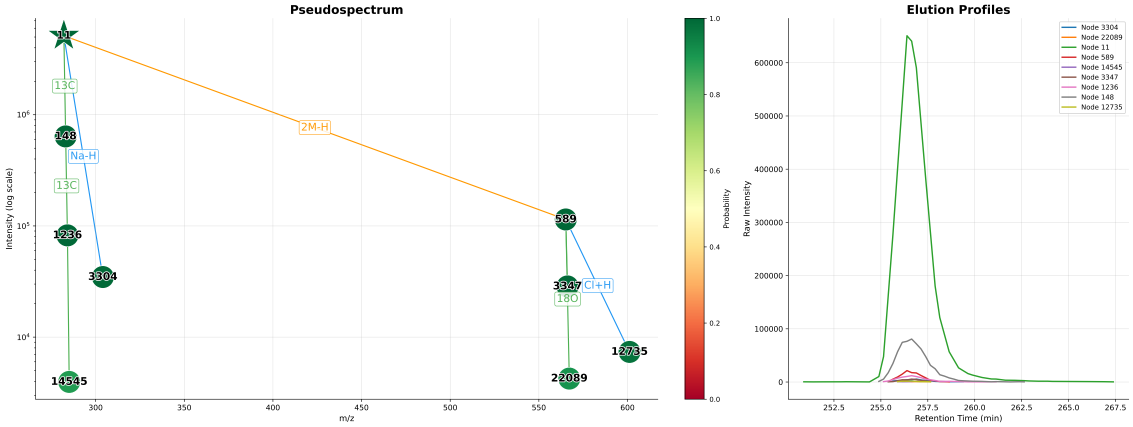


Guanosine

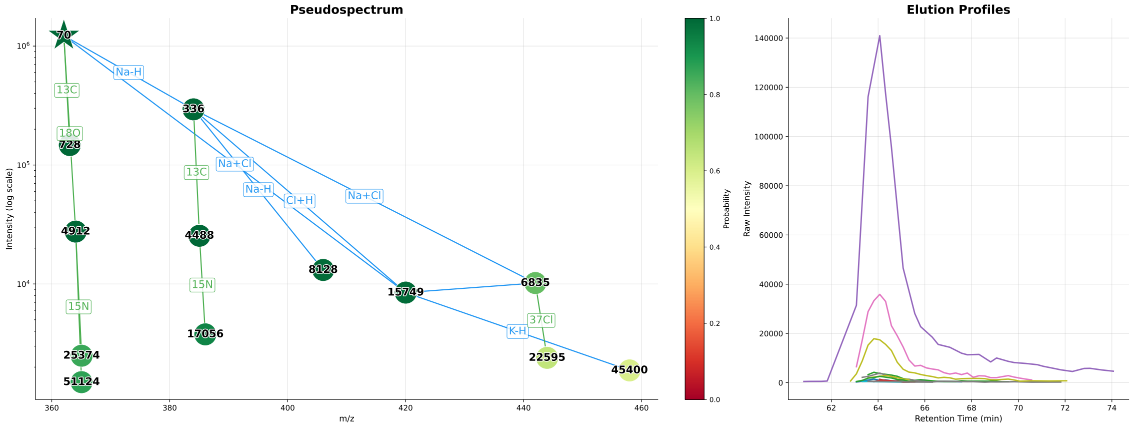


Guanosine monophosphate

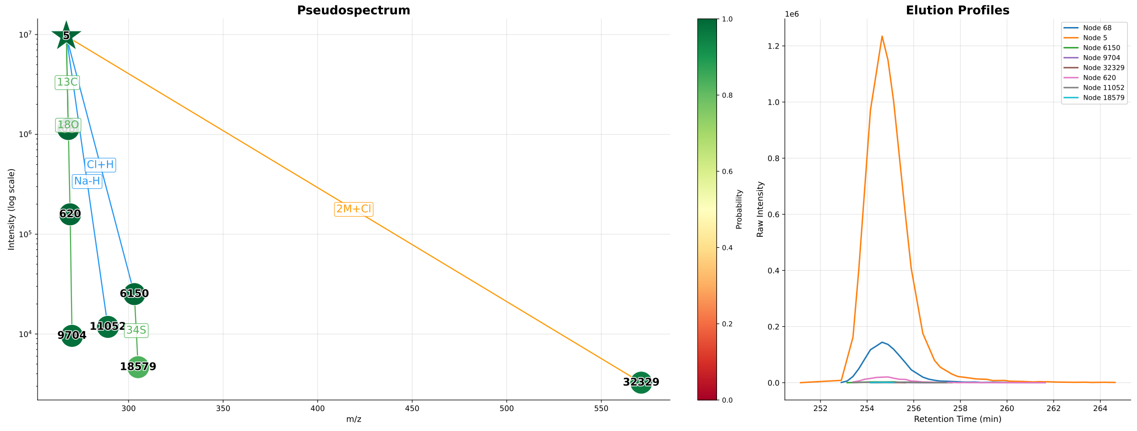


Inosine

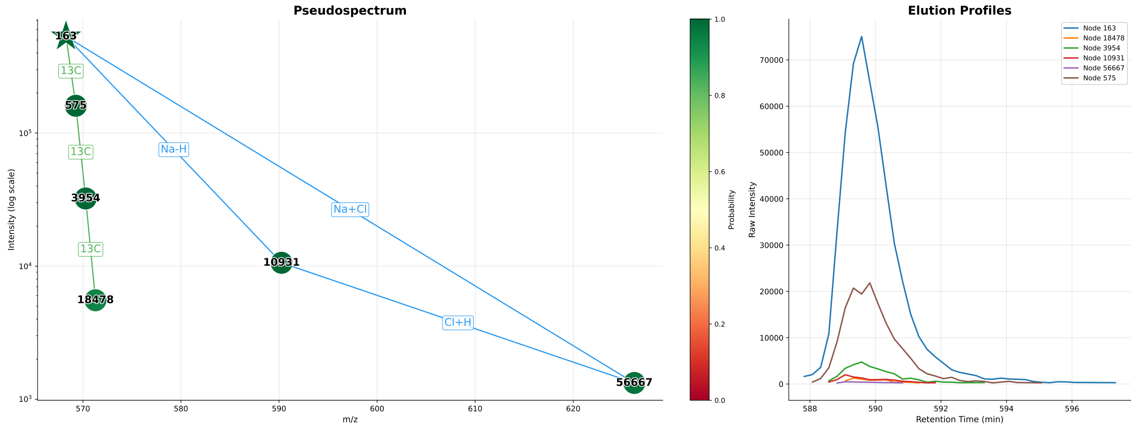


LPSE II

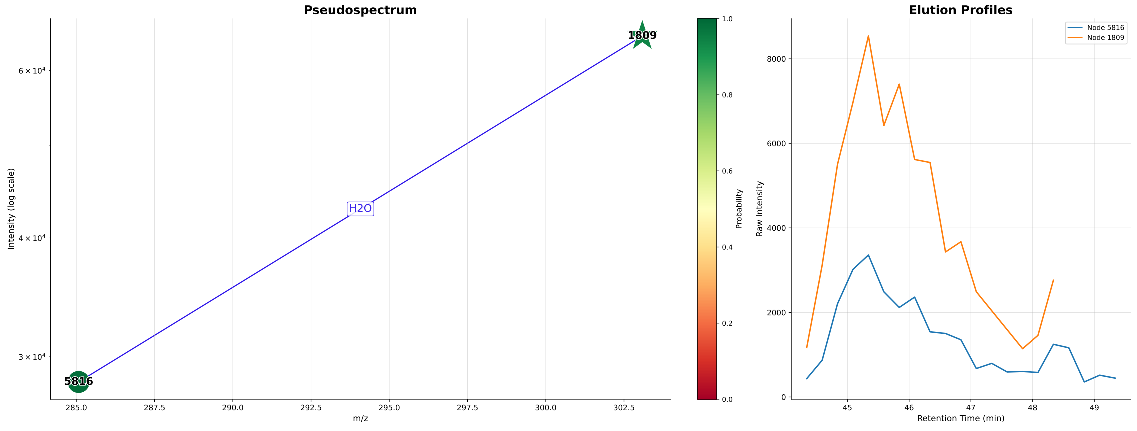


N-Acetylaspartylglutamic acid

**Figure S6:** Re-annotation of Aspartic acid as N-Acetylaspartic acid.

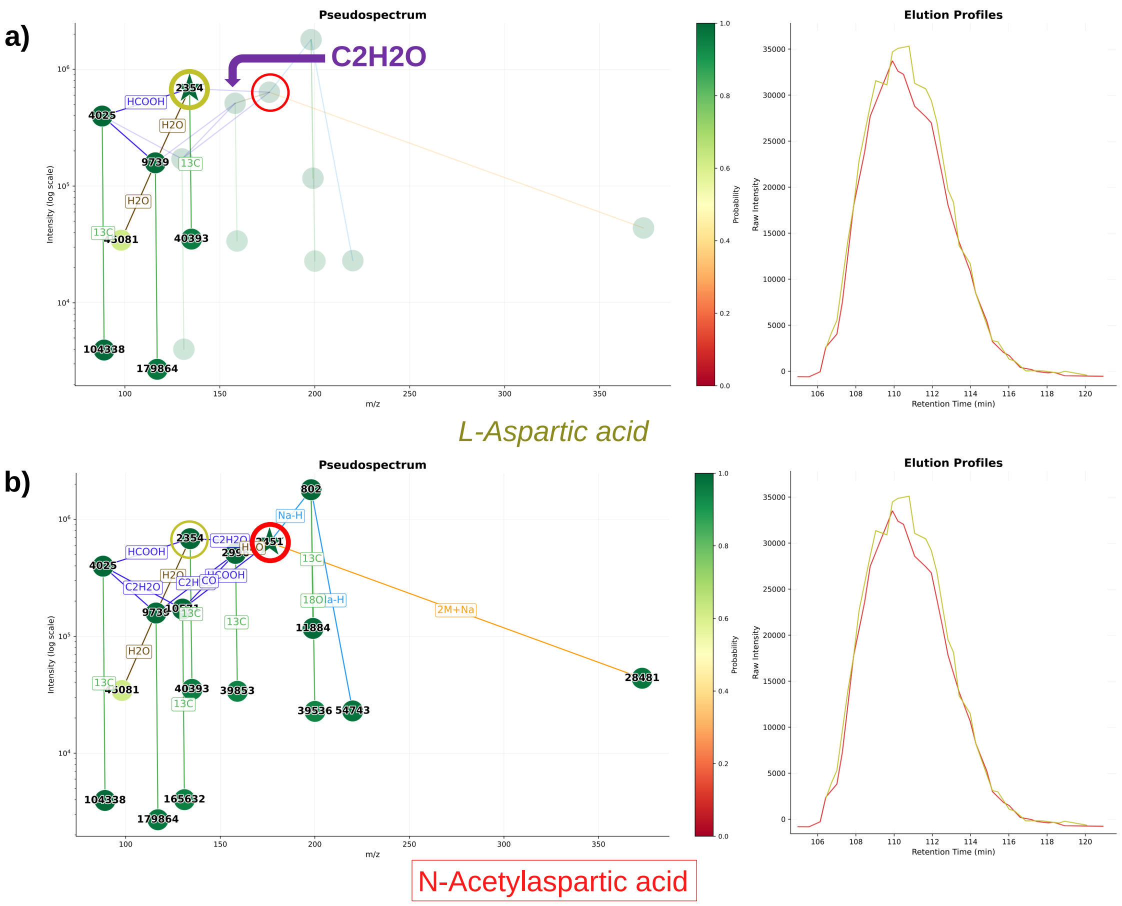


The MassGAT component containing Aspartic acid described in the original benchmark and characterized by the top 2 annotation (**a**) includes a $C_2H_2O$ link with a perfectly correlated peak (red), suggesting that the correct metabolite is in fact N-Acetylaspartic acid (**b**; top 3 annotation).

### Figure S7: MassGAT pseudo-spectra of L-Methionine-oxide in the HIAE cohort.

The MassGAT pseudo-spectra of L-Methionine-oxide in the 18 samples from the MT-BLS103 C18 dataset (Samino et al., 2015) are shown below, for the three groups of controls (CON_BASA) and patients suffering from HyperInsulinaemic Androgen Excess (DIA_BASE, and PIO_BASE). The star corresponds to the targeted ground-truth peak. Nodes (detected ions) are coloured according to their detection probability (green: 1, yellow: 0.5). Edge are labeled by the chemical formula of the m/z difference, and coloured according to type of link (green: isotope, blue: adduct, purple: loss, brown: loss or neutral adduct, orange: dimer or doubly charged).

##### (a) PIO_BASE_603628

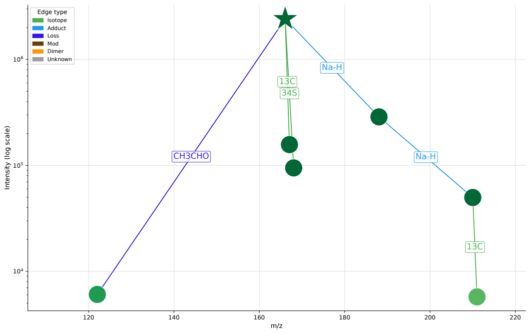


##### (b) PIO_BASE_604520

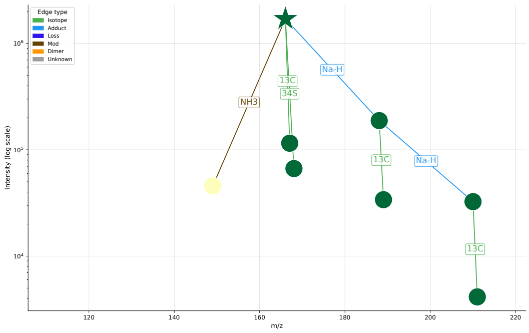


##### (c) PIO_BASE_649358

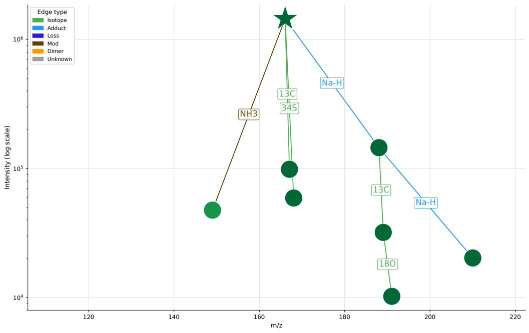


##### (d) PIO_BASE_654139

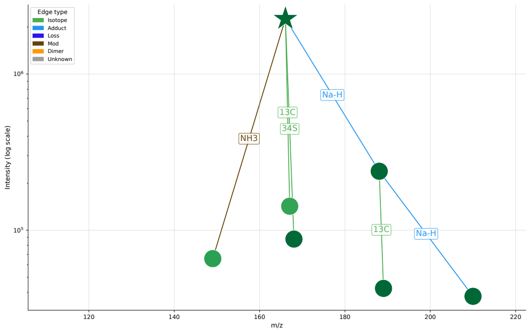


##### (e) PIO_BASE_830049

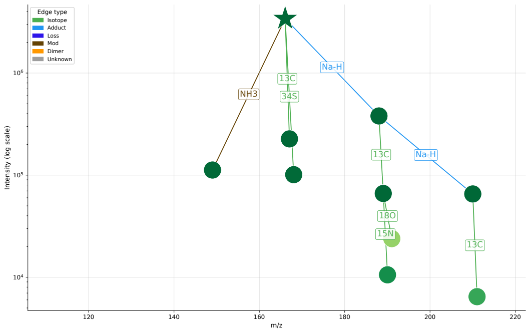

#### (a) CON_BASA_567795

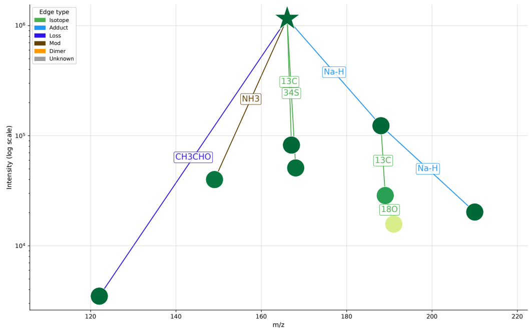


#### (b) CON_BASA_581904

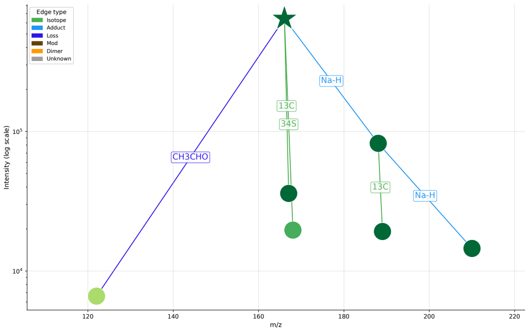


#### (c) CON_BASA_587625

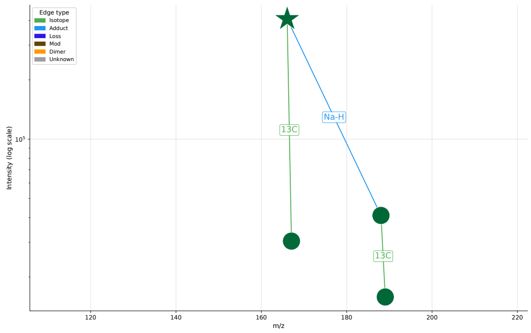


#### (d) CON_BASA_602733

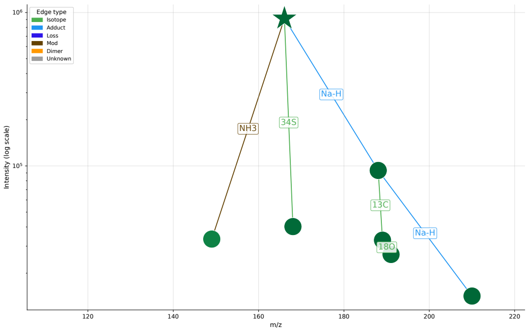


#### (e) CON_BASA_619640

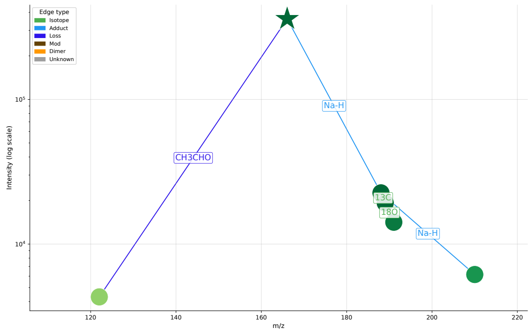


#### (f) CON_BASA_651137

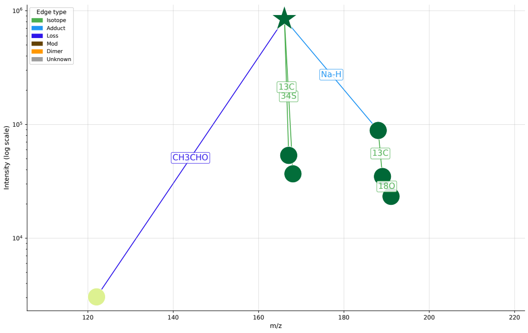


#### (e) CON_BASA_675644

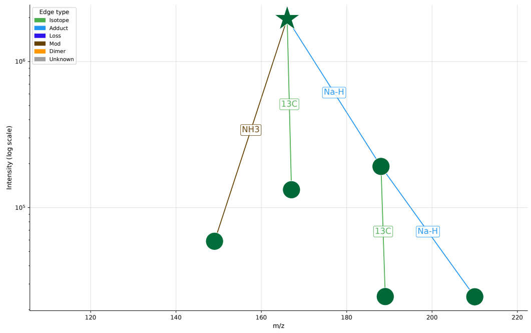


#### (e) CON_BASA_723817

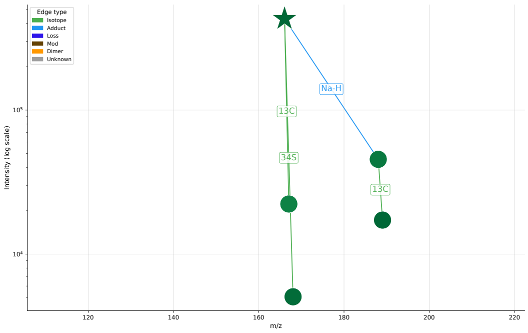

**(a) DIA_BASE_1164621**

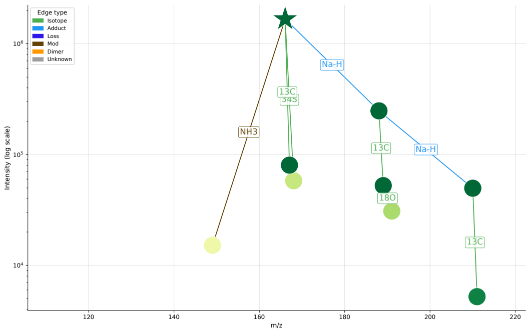


**(b) DIA_BASE_630974**

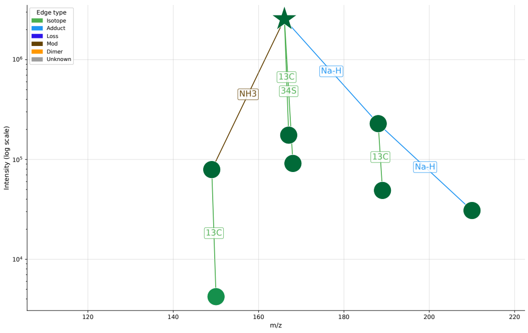


**(c) DIA_BASE_644540**

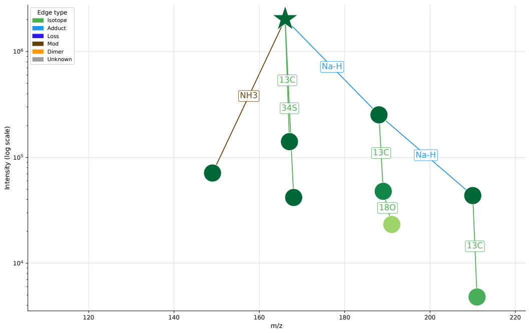


**(d) DIA_BASE_687353**

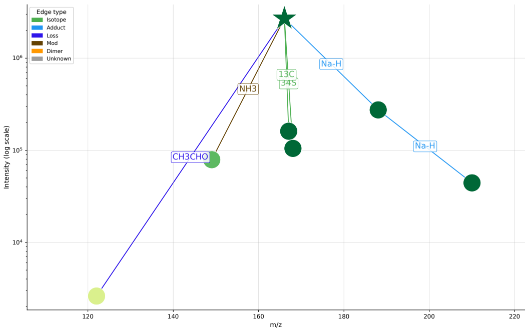


**(e) DIA_BASE_701185**

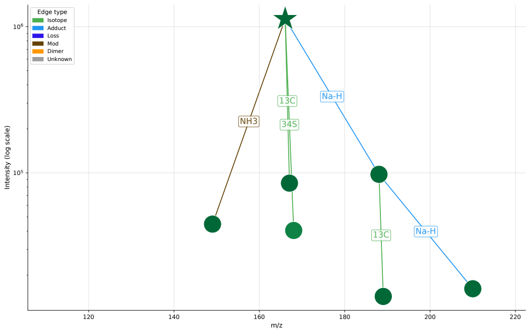

### **Table S1:** List of m/z differences used to build the components.

The table of m/z differences between ion species used by MassGAT to construct the components can be freely modified by the user according to their specific analytical acquisition conditions. The default list (below) contains 32 m/z differences for positive and negative ionisation modes, corresponding to isotopes, adducts, in-source fragments, and dimers. It was compiled from internal data and from the literature as follows.

First, the common m/z differences observed in our in-house spectral data (notably in the database of around 1,000 standards; Roux et al. 2012; Boudah et al. 2014) were calculated and annotated. These data are primarily derived from LC-HRMS acquisitions on Orbitrap instruments (Thermo Fisher Scientific), using Hypersil GOLD C18 (Thermo Fisher Scientific) and Sequant ZIC-pHILIC (Merck) chromatographic columns in positive and negative ionisation modes, respectively. Mobile phases for the C18 column were 100% water in A and 100% acetonitrile in B, both containing 0.1% formic acid. Regarding HILIC, phase A consisted of an aqueous buffer of 10 mM ammonium carbonate in water adjusted to pH 10.5 with ammonium hydroxide, whereas pure acetonitrile was used as solvent B.

Second, this list was compared with the common m/z differences recently identified in 142 public datasets (Nash et al., 2024). Some m/z differences from the literature were thus added (e.g., $CH_2O$, $C_2H_3N$, $C_2H_2O$, $CH_3CHO$), whilst others were not included as they were considered not informative enough in terms of annotation (e.g. multi-charged ions or differences between two distinct fragmentation paths) relative to the risk of increasing the number of accidental relationships.

The final MassGAT list comprises 32 m/z differences, 22 of which are among the 62 most frequent identified by Nash et al. (2024), and further including one isotope ($^{34}S$), two adducts (K-$NH_4$ and $Na_2CO_3$), and six dimer rules (Table S1).

Default m/z differences used by MassGAT to build the components. The *fragment* and *biology* columns refer, respectively, to in-source fragments, and m/z differences naturally occurring between metabolites as a result of biological transformations. The *Nash_freq* column reports the rank among the most frequent m/z differences according to Nash et al. (2024).

| annotation | dmz | ionization | isotope | adduct | fragment | biology | dimer | Nash_freq |
|---|---|---|---|---|---|---|---|---|
| **$^{15}$N** | 0.99703 | both | 1 | | | | | 34 |
| **$^{13}$C** | 1.00335 | both | 1 | | | | | 1 |
| **$^{34}$S** | 1.99579 | both | 1 | | | | | NA |
| **$^{37}$Cl** | 1.99705 | both | 1 | | | | | 3 |
| **$^{18}$O** | 2.00424 | both | 1 | | | | | 25 |
| **2H** | 2.01565 | pos | | | 1 | 1 | | 12 |
| **Na-$NH_4$** | 4.95540 | pos | | 1 | | | | 31 |
| **HCOO-Cl** | 10.02879 | neg | | 1 | | | | 23 |
| **K-Na** | 15.97394 | pos | | 1 | | | | 18 |
| **$NH_3$** | 17.02654 | pos | | 1 | 1 | 1 | | 35 |
| **$H_2O$** | 18.01056 | both | | 1 | 1 | 1 | | 7 |
| **K-$NH_4$** | 20.92934 | pos | | 1 | | | | NA |
| **Na-H** | 21.98194 | both | | 1 | | | | 5 |
| **CO** | 27.99491 | both | | | 1 | 1 | | 32 |
| **$CH_2O$** | 30.01056 | both | | | 1 | 1 | | 37 |
| **Cl+H** | 35.97668 | neg | | 1 | | | | 111 |
| **K-H** | 37.95588 | both | | 1 | | | | 62 |
| **Na+OH** | 39.99250 | neg | | 1 | | | | 54 |
| **$C_2H_3N$** | 41.02654 | both | | 1 | | | | 51 |
| **$C_2H_2O$** | 42.01056 | pos | | | 1 | 1 | | 46 |
| **$CO_2$** | 43.98982 | both | | | 1 | 1 | | 52 |
| **$CH_3CHO$** | 44.02621 | both | | | 1 | 1 | | 8 |
| **HCOOH** | 46.00547 | pos | | | 1 | 1 | | 17 |
| **H+HCOO** | 46.00547 | neg | | 1 | | 1 | | 17 |
| **Na+Cl** | 57.95862 | neg | | 1 | | | | 16 |
| **HCOO+Na** | 67.98741 | neg | | 1 | | | | 2 |
| **$Na_2CO_3$** | 105.96426 | neg | | 1 | | | | NA |
| **2M-H** | -1.00783 | neg | | | | | 1 | NA |
| **2M+H** | 1.00783 | pos | | | | | 1 | NA |
| **2M+$NH_4$** | 18.03382 | pos | | | | | 1 | NA |
| **2M+Na** | 22.98977 | pos | | | | | 1 | NA |
| **2M+Cl** | 34.96885 | neg | | | | | 1 | NA |
| **2M+HCOO** | 44.99765 | neg | | | | | 1 | NA |

#### **Table S2:** Parameter values used to benchmark the data processing software (TripleTOF 6600 and QE HF datasets).

The parameters used for XCMS (respectively, 3D-MSNet) are those reported in the initial benchmark by Li et al., 2018 (respectively, those reported in the original article by Wang et al., 2023). For MassGAT, the minimum value of 8 scans per ion trace was used (instead of the default 4 scans) to account for the high frequency of acquisition (5Hz) in these datasets.

| **Software** | **Parameter** | **TripleTOF** | **QE HF** |
|---|---|---|---|
| **XCMS** | algorithm | centWave | centWave |
| | mzdiff | 0.01 | 0.005 |
| | mzwid | 0.015 | 0.005 |
| | prefilter | 10 | 100 |
| | bw | 5 | 5 |
| | minfrac | 0.5 | 0.5 |
| | noise | 0 | 0 |
| **asari** | mode | pos | pos |
| | mz_tol. (ppm) | 10 | 5 |
| | min_intens. thresh. | 100 | 100 |
| | autoheight | True | True |
| **3D-MSNet** | *Point cloud extraction* | | |
| | window_mz_width | 0.8 | 0.4 |
| | window_rt_width | 6 | 6 |
| | min_intensity | 128 | 10000 |
| | from_mz | 100 | 100 |
| | to_mz | 1300 | 1300 |
| | from_rt | 0 | 0 |
| | to_rt | 40 | 40 |
| | max_peak_mz_width | 0.1 | 0.05 |
| | max_peak_rt_width | 1 | 1 |
| | *Feature detection* | | |
| | experiment | msnet_20220215 | |
| | epoch | 300 | 300 |
| | mass_analyzer | tof | orbitrap |
| | mz_resolution | 35000 | 60000 |
| | resolution_mz | 956 | 200 |
| | rt_fwhm | 0.1 | 0.08 |
| | center_threshold | 0.5 | 0.5 |
| | block_rt_width | 6 | 6 |
| | block_mz_width | 0.8 | 0.4 |
| **MassGAT** | *ionshed* | | |
| | rttol | 5.0 | 5.0 |
| | ppm | 3 | 3 |
| | mz_res | 0.001 | 0.001 |
| | rt_res | 0.1 | 0.1 |
| | *Graph Construction* | | |
| | rt_window | 5.0 | 5.0 |
| | mz_tolerance | 0.001 | 0.001 |
| | isotope_corr_thresh. | 0.6 | 0.6 |
| | adduct_corr_thresh. | 0.85 | 0.85 |
| | dimer_corr_thresh. | 0.85 | 0.85 |
| | *Model* | | |
| | hidden_dim | 32 | 32 |
| | embedding_dim | 64 | 64 |

#### **Table S3:** Annotations from CliqueMS and MassGAT on the IRS2-KO sample (positive ionization).

The visualizations corresponding to the following annotated pseudo-spectra are shown on Fig S4.

| **Metabolite** | **Annotation** | **CliqueMS** | | **MassGAT** | |
|---|---|---|---|---|---|
| | | **Isotopes** | **Rank** | **Isotopes** | **Rank** |
| 2-Methylbutyroylcarnitine | (M+H)+ | - | - | 2 | 1 |
| 2-Pyrrolidinone | (M+2Na+H2O-H)+ | - | - | 0 | 1 |
| - | (M+H)+ | - | - | 0 | 1 |
| | (M+H+H2O)+ | - | - | 1 | 1 |
| | (M+H+H2O-NH3)+ | - | - | 1 | 1 |
| | (M+Na+H2O)+ | - | - | 0 | 1 |
| Adenine | (M+H)+ | 1 | 1 | 3 | 1 |
| - | (M+H-NH3)+ | 1 | 1 | 1 | 1 |
| Adenylosuccinic acid | (M+H)+ | - | - | 2 | 1 |
| GABA | (M+H)+ | 1 | 1 | 1 | 3 |
| - | (M+H-H2O)+ | 0 | 1 | 0 | 3 |
| | (M+H-NH3)+ | 0 | 1 | 1 | 3 |
| | (M+Na)+ | 0 | 1 | 0 | 3 |
| | (M+2Na-H)+ | - | - | 0 | 3 |
| Glutathione | (M+H)+ | 1 | 1 | 3 | 1 |
| - | (M+Na)+ | 0 | 1 | 1 | 1 |
| Guanine | (M+H)+ | 2 | 1 | 2 | 2 |
| - | (M+H-H2O)+ | 0 | 1 | 0 | 2 |
| | (M+H-NH3)+ | 1 | 1 | 1 | 2 |
| | (M+H+H2O-2NH3)+ | - | - | 0 | 2 |
| | (M+H+H2O-CO2-NH3)+ | - | - | 0 | 2 |
| | (M+H+H2O-NH3)+ | - | - | 0 | 2 |
| Guanosine | (2M+H)+ | 1 | 1 | 2 | 1 |
| - | (M+H)+ | 3 | 1 | 3 | 1 |
| | (M+Na)+ | 0 | 1 | 1 | 1 |
| Inosine | (2M+H)+ | 1 | 1 | 3 | 1 |
| - | (2M+Na)+ | 2 | 1 | 2 | 1 |
| | (M+H)+ | 2 | 1 | 2 | 1 |
| | (M+K)+ | 1 | 1 | 0 | 1 |
| | (M+Na)+ | 1 | 1 | 2 | 1 |
| | (M+H-H2O)+ | 0 | 1 | - | - |
| | (2M+K)+ | - | - | 0 | 1 |
| L-2-Aminoadipic acid | (M+H)+ | 0 | 1 | 0 | 1 |
| - | (M+H-H2O)+ | 0 | 1 | 0 | 1 |
| | (M+H-CO)+ | 1 | 1 | - | - |
| | (M+H-HCOOH)+ | 1 | 1 | - | - |
| L-Alanine | (M+H)+ | - | - | 1 | 1 |
| L-Ascorbic acid | (M+H)+ | 0 | 2 | 3 | 3 |
| - | (M+Na)+ | 0 | 2 | 1 | 3 |
| | (M+H-HCOOH)+ | 0 | 2 | - | - |
| | (M+H-CO2)+ | - | - | 0 | 3 |
| | (M+H-CO2-2H2O)+ | - | - | 1 | 3 |
| | (M+H-CO2-H2O)+ | - | - | 0 | 3 |
| | (M+H-CO2-HCOOH)+ | - | - | 0 | 3 |
| | (M-3H-CO-CO2)+ | - | - | 0 | 3 |
| | (M-3H-CO2)+ | - | - | 0 | 3 |
| | (M-3H-CO2-H2O)+ | - | - | 0 | 3 |
| | (M-H-CO2-2H2O)+ | - | - | 1 | 3 |
| | (M-H-CO2-H2O)+ | - | - | 1 | 3 |
| L-Aspartic acid | (M+H)+ | 1 | 2 | 1 | 4 |
| - | (M+H-H2O)+ | 1 | 2 | 1 | 4 |
| | (2M+Na)+ | 1 | 2 | - | - |
| | (M+H+HCOONa)+ | 2 | 2 | - | - |
| | (M+H-C2H2O)+ | 0 | 2 | - | - |
| | (M+H-2H2O)+ | - | - | 0 | 4 |
| | (M+H-HCOOH)+ | - | - | 1 | 4 |
| L-Glutamic acid | (2M+Na)+ | 0 | 1 | 0 | 1 |

*Continued on next page*

| Metabolite | Annotation | CliqueMS | | MassGAT | |
|---|---|---|---|---|---|
| | | Isotopes | Rank | Isotopes | Rank |
| - | (M+H)+ | 2 | 1 | 3 | 1 |
| | (M+H-H2O)+ | 1 | 1 | 2 | 1 |
| | (M+Na)+ | 1 | 1 | 2 | 1 |
| | (2M+H)+ | 0 | 1 | - | - |
| | (M+H+HCOONa)+ | 2 | 1 | - | - |
| | (M+H-NH3)+ | 0 | 1 | - | - |
| | (M+K)+ | 0 | 1 | - | - |
| | (2M+2Na-H)+ | - | - | 0 | 1 |
| | (2M+3Na-2H)+ | - | - | 1 | 1 |
| | (2M+4Na-3H)+ | - | - | 0 | 1 |
| | (2M+5Na-4H)+ | - | - | 1 | 1 |
| | (M+2Na+K-2H)+ | - | - | 0 | 1 |
| | (M+2Na-H)+ | - | - | 3 | 1 |
| | (M+2Na-H-H2O)+ | - | - | 0 | 1 |
| | (M+2Na-NH4)+ | - | - | 0 | 1 |
| | (M+3Na-2H)+ | - | - | 1 | 1 |
| | (M+H-H2O-HCOOH)+ | - | - | 1 | 1 |
| | (M+H-HCOOH)+ | - | - | 1 | 1 |
| | (M+H-HCOOH-NH3)+ | - | - | 0 | 1 |
| | (M+Na+K-H)+ | - | - | 0 | 1 |
| | (M+Na-H2O)+ | - | - | 2 | 1 |
| NAD | (M+2H)2+ | 2 | 1 | 1 | 2 |
| - | (M+H)+ | 3 | 1 | 3 | 2 |
| | (M+Na)+ | 0 | 1 | - | - |
| Oxigluthatione | (M+2H)2+ | 1 | 1 | 0 | 1 |
| - | (M+H)+ | 3 | 1 | 5 | 1 |
| | (M+H-H2O)+ | 2 | 1 | 0 | 1 |
| | (2M+H)+ | 2 | 1 | - | - |
| | (M+H+HCOONa)+ | 1 | 1 | - | - |
| | (M+H+Na2CO3)+ | 0 | 1 | - | - |
| | (M+H+NaOH)+ | 0 | 1 | - | - |
| | (M+H-C2H2O)+ | 1 | 1 | - | - |
| | (M+H-CH2O)+ | 1 | 1 | - | - |
| | (M+H-CH3CHO)+ | 1 | 1 | - | - |
| | (M+H-NH3)+ | 2 | 1 | - | - |
| | (M+K)+ | 0 | 1 | - | - |
| | (M+Na)+ | 1 | 1 | - | - |
| | (M+2H-H2O)2+ | - | - | 0 | 1 |
| | (M+2H-NH3)2+ | - | - | 0 | 1 |
| PC | (M+H)+ | 2 | 1 | 2 | 1 |
| - | (M+K)+ | 0 | 1 | 0 | 1 |
| | (M+Na)+ | 1 | 1 | 2 | 1 |
| Taurine | (2M+H)+ | 0 | 1 | 0 | 1 |
| - | (M+H)+ | 2 | 1 | 2 | 1 |
| | (M+H-H2O)+ | 0 | 1 | 1 | 1 |
| | (M+Na)+ | 1 | 1 | 2 | 1 |
| | (2M+Na)+ | 0 | 1 | - | - |
| | (2M+2Na+K-2H)+ | - | - | 0 | 1 |
| | (2M+2Na-H)+ | - | - | 0 | 1 |
| | (2M+3Na-2H)+ | - | - | 1 | 1 |
| | (M+2Na-H)+ | - | - | 2 | 1 |
| | (M+Na+K-H)+ | - | - | 2 | 1 |
| Uracil | (M+H)+ | 1 | 1 | 2 | 1 |
| - | (M+H-H2O)+ | 0 | 1 | 0 | 1 |
| | (M+H-NH3)+ | 0 | 1 | 0 | 1 |
| Xanthine | (M+H)+ | 1 | 1 | 2 | 1 |
| - | (M+H-NH3)+ | 0 | 1 | 0 | 1 |
| | (M+Na)+ | 0 | 1 | 0 | 1 |
| | (M+H+2H2O-CO2-NH3)+ | - | - | 0 | 1 |
| | (M+H+H2O-CO2-NH3)+ | - | - | 1 | 1 |
| | (M+H+H2O-NH3)+ | - | - | 0 | 1 |
| | (M+H-H2O)+ | - | - | 0 | 1 |

MassGAT componentization parameters were configured with a 0.0015 $m/z$ tolerance, an isotope threshold of 0.5, and an adduct/dimer threshold of 0.85.

#### **Table S4:** Annotations from CliqueMS and MassGAT on the IRS2-KO deficient mice (negative ionization).

The visualizations corresponding to the following annotated pseudo-spectra are shown on Fig S5.

| Metabolite | Annotation | CliqueMS | | MassGAT | |
|---|---|---|---|---|---|
| | | Isotopes | Rank | Isotopes | Rank |
| 2-hydroxy-2-methyl-butyric acid | (M-H)- | - | - | 1 | 1 |
| Adenine | (M-H)- | - | - | 1 | 1 |
| Adenosine | (2M-H)- | 2 | 1 | 2 | 1 |
| - | (M+Cl)- | 1 | 1 | 3 | 1 |
| | (M+HCOO)- | 2 | 1 | 1 | 1 |
| | (M-H)- | 2 | 1 | 2 | 1 |
| | (M-H-CH2O)- | 0 | 1 | - | - |
| | (M-H-CH3CHO)- | 0 | 1 | - | - |
| | (M-H-H2O)- | 1 | 1 | - | - |
| | (M+Na+2HCOO)- | - | - | 3 | 1 |
| Ascorbic acid | - | - | - | - | - |
| FAD | (M-H)- | - | - | 0 | 1 |
| Guanosine | (2M+Cl)- | 0 | 1 | 0 | 1 |
| - | (2M-H)- | 2 | 1 | 2 | 1 |
| | (M-H)- | 3 | 1 | 3 | 1 |
| | (M+Cl)- | 0 | 1 | - | - |
| | (M-2H+K)- | 0 | 1 | - | - |
| | (M-2H+Na)- | 0 | 1 | - | - |
| | (M+Na-2H)- | - | - | 0 | 1 |
| Guanosine monophosphate | (M-H)- | 2 | 2 | 4 | 1 |
| - | (M-2H+Na)- | 1 | 2 | - | - |
| | (M+2Na+Cl-2H)- | - | - | 1 | 1 |
| | (M+2Na-3H)- | - | - | 0 | 1 |
| | (M+Na+Cl-H)- | - | - | 0 | 1 |
| | (M+Na+K+Cl-2H)- | - | - | 0 | 1 |
| | (M+Na-2H)- | - | - | 2 | 1 |
| Hypoxanthine | (M+Na-2H)- | - | - | 0 | 1 |
| - | (M-H)- | - | - | 2 | 1 |
| Inosine | (M+Cl)- | 0 | 1 | 1 | 1 |
| - | (M-H)- | 3 | 1 | 3 | 1 |
| | (2M-H)- | 2 | 1 | - | - |
| | (M-2H+K)- | 0 | 1 | - | - |
| | (M-2H+Na)- | 0 | 1 | - | - |
| | (2M+Cl)- | - | - | 0 | 1 |
| | (M+Na-2H)- | - | - | 0 | 1 |
| L-Glutamic acid | (M-H)- | - | - | 2 | 3 |
| - | (M-H-H2O)- | - | - | 1 | 3 |
| LPI I | (M-H)- | - | - | 3 | 1 |
| LPI II | (M-H)- | - | - | 3 | 1 |
| LPI III | (M-H)- | - | - | 3 | 1 |
| LPSE I | (M-H)- | - | - | 2 | 1 |
| LPSE II | (M-H)- | 2 | 2 | 3 | 1 |
| - | (M+HCOO)- | 1 | 2 | - | - |
| | (M-2H+Na)- | 0 | 2 | - | - |
| | (M-H-NH3)- | 1 | 2 | - | - |
| | (M+Na+Cl-H)- | - | - | 0 | 1 |
| | (M+Na-2H)- | - | - | 0 | 1 |
| N-Acetylaspartylglutamic acid | (M-H)- | 1 | 1 | 0 | 1 |
| - | (M-H-H2O)- | 0 | 1 | 0 | 1 |
| S-Adenosylhomocysteine | (M-H)- | - | - | 2 | 1 |
| Taurine | (M+2Na+2Cl-H)- | - | - | 2 | 1 |
| - | (M+3Na+3Cl-H)- | - | - | 2 | 1 |
| | (M+4Na+4Cl-H)- | - | - | 0 | 1 |
| | (M+Na+Cl-H)- | - | - | 4 | 1 |
| | (M-H)- | - | - | 3 | 1 |

MassGAT componentization parameters were configured with a 0.0015 $m/z$ tolerance, an isotope threshold of 0.5, and an adduct/dimer threshold of 0.8.

**Table S5:** Annotations from CliqueMS and MassGAT on the HIAE cohort (MTBLS103 dataset).

| Metabolite | Annotation | CliqueMS | | | MassGAT | | |
|---|---|---|---|---|---|---|---|
| | | Samples | Rank | Isotopes | Samples | Rank | Isotopes |
| Asp-Phe | (M+H)+ | 5/18 | 1 | 3 | 18/18 | 1 | 2 |
| | (M+H-H2O)+ | 2/18 | 1 | 1 | 12/18 | 1 | 1 |
| | (M+Na)+ | 4/18 | 1 | 1 | - | - | - |
| Choline | (M+H)+ | - | - | - | 12/18 | 1 | 3 |
| Glu-Glu | (M+H)+ | 14/18 | 1 | 1.5 | 16/18 | 1 | 3 |
| | (M+H-H2O)+ | 10/18 | 1 | 1 | 16/18 | 1 | 2 |
| | (M+Na)+ | 14/18 | 1 | 1 | 10/18 | 1 | 1 |
| | (M+H-2H2O)+ | - | - | - | 3/18 | 1 | 1 |
| | (M+H-CO2)+ | - | - | - | 3/18 | 2 | 1 |
| Glutamate | (M+H)+ | 17/18 | 1 | 3 | 9/18 | 2 | 3 |
| | (M+Na)+ | 14/18 | 1 | 3 | 9/18 | 2 | 2 |
| | (M+H-NH3)+ | - | - | - | 8/18 | 2 | 1 |
| | (M+Na-H2O)+ | 8/18 | 1 | 2 | 9/18 | 2 | 2 |
| | (M+2Na-H)+ | - | - | - | 9/18 | 2 | 1 |
| | (M+3Na-2H)+ | - | - | - | 6/18 | 2 | 1 |
| | (M+H-HCOOH)+ | - | - | - | 9/18 | 2 | 2 |
| | (M+H-CO-NH3)+ | - | - | - | 5/18 | 2 | 1 |
| | (M+K)+ | 4/18 | 1 | 1 | - | - | - |
| | (M+Na+NH3-H2O)+ | - | - | - | 6/18 | 2.5 | 1 |
| | (2M+H)+ | 8/18 | 1 | 1 | 3/18 | 2 | 1 |
| | (M+H-H2O)+ | 1/18 | 1 | 1 | - | - | - |
| | (M-H+2Na)+ | 16/18 | 1 | 1 | - | - | - |
| | (M+H+H2O-HCOOH)+ | - | - | - | 9/18 | 2 | 1 |
| | (M+2Na-H-CO)+ | - | - | - | 9/18 | 2 | 1 |
| | (M+Na+H2O-HCOOH)+ | - | - | - | 7/18 | 2 | 1 |
| | (M+Na+NH3-2H2O)+ | - | - | - | 5/18 | 2 | 1 |
| | (M+H-H2O-NH3)+ | - | - | - | 4/18 | 2 | 1 |
| | (M+2Na+NH3-H-H2O)+ | - | - | - | 3/18 | 2 | 1 |
| Glutamine | (M+H)+ | 14/18 | 1 | 2 | 11/18 | 1 | 1 |
| | (M+H-NH3)+ | 14/18 | 1 | 1 | 10/18 | 1 | 2 |
| | (M+H-HCOOH)+ | - | - | - | 10/18 | 1 | 1 |
| Glutamyl-Taurine | (M+H)+ | 15/18 | 1 | 2 | 16/18 | 1 | 1 |
| | (M+Na)+ | 9/18 | 1 | 2 | 11/18 | 1 | 2 |
| | (M+2Na-H)+ | - | - | - | 10/18 | 1 | 1 |
| | (M-H+2Na)+ | 15/18 | 1 | 1 | - | - | - |
| L-Methionine S-oxide | (M+H)+ | 18/18 | 1 | 2 | 18/18 | 1 | 3 |
| | (M+Na)+ | 18/18 | 1 | 1 | 18/18 | 1 | 2 |
| | (M+2Na-H)+ | - | - | - | 16/18 | 1 | 1 |
| | (M+H-NH3)+ | 14/18 | 1 | 1.5 | 11/18 | 1 | 1 |
| | (M+H-CH3CHO)+ | - | - | - | 7/18 | 1 | 1 |
| | (M-H+2Na)+ | 18/18 | 1 | 2 | - | - | - |
| Val Glu | (M+H)+ | 3/18 | 2 | 3 | 18/18 | 1 | 3 |
| | (M+H-NH3)+ | 3/18 | 2 | 1 | 5/18 | 1 | 1 |
| | (M+Na)+ | - | - | - | 3/18 | 1 | 1 |
| g-D-Glutamylglycine | (M+H)+ | 18/18 | 1 | 3 | 15/18 | 1 | 3 |
| | (M+Na)+ | 17/18 | 1 | 1 | 15/18 | 1 | 2 |
| | (M+2Na-H)+ | - | - | - | 13/18 | 1 | 1 |
| | (M+K)+ | 1/18 | 2 | 1 | - | - | - |
| | (M-H+2Na)+ | 4/18 | 1 | 1 | - | - | - |

C18 column (18 samples).

| Metabolite | Annotation | CliqueMS | | | MassGAT | | |
|---|---|---|---|---|---|---|---|
| | | Samples | Rank | Isotopes | Samples | Rank | Isotopes |
| 5-oxoproline | (M+H)+ | 11/13 | 2 | 1 | 11/13 | 2 | 1 |
| | (M+H-HCOOH)+ | - | - | - | 4/13 | 2 | 0 |
| | (M+Na)+ | - | - | - | 3/13 | 2 | 0 |
| | (M+2Na-H)+ | - | - | - | 3/13 | 2 | 1 |
| | (M-2H+3Na)+ | 10/13 | 2 | 1 | - | - | - |
| | (M-H+2Na)+ | 11/13 | 2 | 2 | - | - | - |
| | (2M+3Na-2H)+ | - | - | - | 3/13 | 2 | 0 |
| | (2M+4Na-3H)+ | - | - | - | 3/13 | 2 | 0 |
| | (2M+3Na-2H-CH3CHO)+ | - | - | - | 3/13 | 2 | 0 |
| Choline | (M+H)+ | 7/13 | 1 | 3 | 10/13 | 1 | 0 |
| | (2M+H)+ | 7/13 | 1 | 2 | - | - | - |
| Glutamate | (M+H)+ | 10/13 | 1 | 2 | 12/13 | 1 | 2 |
| | (M+Na)+ | 8/13 | 1 | 1 | 12/13 | 1 | 0 |
| | (M+2Na-H)+ | - | - | - | 12/13 | 1 | 1 |
| | (M+3Na-2H)+ | - | - | - | 12/13 | 1 | 1 |
| | (M+H-HCOOH)+ | - | - | - | 11/13 | 1 | 1 |
| | (M+Na+K-H)+ | - | - | - | 10/13 | 1 | 0 |
| | (M+H-H2O-HCOOH)+ | - | - | - | 8/13 | 1 | 0 |
| | (M+2Na-H-H2O)+ | - | - | - | 3/13 | 1 | 0 |
| | (M+2Na-NH4)+ | - | - | - | 7/13 | 1 | 0 |
| | (M+H-H2O)+ | - | - | - | 4/13 | 1 | 1 |
| | (M+2Na+K-2H)+ | - | - | - | 4/13 | 1 | 0 |
| | (M+H+C2H3N-H2O-HCOOH)+ | - | - | - | 3/13 | 2 | 0 |
| | (M+H-NH3)+ | 5/13 | 1 | 1 | - | - | - |
| | (M-2H+3Na)+ | 8/13 | 1 | 2 | - | - | - |
| | (M-H+2Na)+ | 6/13 | 1.5 | 3 | - | - | - |
| | (M+3Na+C2H3N-2H)+ | - | - | - | 3/13 | 2 | 0 |
| | (2M+3Na-2H)+ | - | - | - | 9/13 | 1 | 0 |
| | (2M+4Na-3H)+ | - | - | - | 12/13 | 1 | 0 |
| | (2M+5Na-4H)+ | - | - | - | 12/13 | 1 | 0 |
| | (2M+4Na-3H-C2H2O)+ | - | - | - | 5/13 | 1 | 0 |
| | (2M+5Na-4H-C2H2O)+ | - | - | - | 4/13 | 2.5 | 0 |
| | (2M+5Na-4H-CH3CHO)+ | - | - | - | 3/13 | 1 | 0 |
| | (2M+4Na-3H-HCOOH)+ | - | - | - | 3/13 | 1 | 0 |
| L-Methionine S-oxide | (M+H)+ | 4/13 | 1 | 1.5 | 13/13 | 1 | 1 |
| | (M+Na)+ | 3/13 | 1 | 1 | 6/13 | 1 | 0 |
| | (M+2Na-H)+ | - | - | - | 6/13 | 1 | 0 |
| | (M-H+2Na)+ | 2/13 | 1 | 1 | - | - | - |
| Methionine | (M+H)+ | 12/13 | 1 | 2 | 13/13 | 1 | 1 |
| | (M+H-NH3)+ | - | - | - | 12/13 | 1 | 0 |
| | (M+H-HCOOH)+ | - | - | - | 6/13 | 1 | 0 |
| | (M+Na)+ | 2/13 | 1 | 1 | - | - | - |
| | (M-H+2Na)+ | 2/13 | 1 | 2 | - | - | - |
| | (M-H+NH3)+ | 12/13 | 1 | 1 | - | - | - |
| Taurine | (M+H)+ | 12/13 | 1.5 | 2 | 13/13 | 1 | 2 |
| | (M+Na)+ | 12/13 | 2 | 1 | 13/13 | 1 | 0 |
| | (M+H-H2O)+ | 12/13 | 1 | 1 | 13/13 | 1 | 0 |
| | (M+H+NH3)+ | - | - | - | 12/13 | 1 | 0 |
| | (M+2Na-H)+ | - | - | - | 11/13 | 1 | 0 |
| | (M+K)+ | 2/13 | 1 | 1 | - | - | - |
| | (M+NH4)+ | 12/13 | 1 | 1 | - | - | - |
| | (M-H+2Na)+ | 11/13 | 1 | 1 | - | - | - |
| | (2M+H)+ | 12/13 | 2 | 2 | 13/13 | 1 | 2 |
| | (2M+Na)+ | 8/13 | 2 | 1 | 4/13 | 1 | 0 |
| | (2M+2Na-H)+ | - | - | - | 4/13 | 1 | 0 |

HILIC column (13 samples).

### References

S. H. Bach, M. Broecheler, B. Huang, and L. Getoor. Hinge-loss markov random fields and probabilistic soft logic. *Journal of Machine Learning Research*, 18(109):1–67, 2017.

S. Boudah, M.-F. Olivier, S. Aros-Calt, L. Oliveira, F. Fenaille, J.-C. Tabet, and C. Junot. Annotation of the human serum metabolome by coupling three liquid chromatography methods to high-resolution mass spectrometry. *Journal of Chromatography B*, 966: 34–47, Sept. 2014. ISSN 1570-0232. doi: 10.1016/j.jchromb.2014.04.025.

X. Bresson and T. Laurent. Residual gated graph convnets. *arXiv preprint arXiv:1711.07553*, 2017.

B. C. DeFelice, S. S. Mehta, S. Samra, T. Cajka, B. Wancewicz, J. F. Fahrmann, and O. Fiehn. Mass spectral feature list optimizer (MS-FLO): A tool to minimize false positive peak reports in untargeted liquid chromatography-mass spectroscopy (LC-MS) data processing. *Analytical Chemistry*, 89(6):3250–3255, 2017. ISSN 0003-2700. doi: 10.1021/acs.analchem.6b04372.

M. Fey and J. E. Lenssen. Fast graph representation learning with pytorch geometric. *arXiv preprint arXiv:1903.02428*, 2019.

E. Giunchiglia and T. Lukasiewicz. Coherent hierarchical multi-label classification networks. *Advances in neural information processing systems*, 33:9662–9673, 2020.

D. P. Kingma and J. Ba. Adam: A method for stochastic optimization. *arXiv preprint arXiv:1412.6980*, 2014.

C. Kuhl, R. Tautenhahn, C. Böttcher, T. R. Larson, and S. Neumann. CAMERA: An Integrated Strategy for Compound Spectra Extraction and Annotation of Liquid Chromatography/Mass Spectrometry Data Sets. *Analytical Chemistry*, 84(1):283–289, Jan. 2012. ISSN 0003-2700. doi: 10.1021/ac202450g.

S. Li, A. Siddiqa, M. Thapa, Y. Chi, and S. Zheng. Trackable and scalable LC-MS metabolomics data processing using asari. *Nature Communications*, 14(1):4113, July 2023. ISSN 2041-1723. doi: 10.1038/s41467-023-39889-1.

Z. Li, Y. Lu, Y. Guo, H. Cao, Q. Wang, and W. Shui. Comprehensive evaluation of untargeted metabolomics data processing software in feature detection, quantification and discriminating marker selection. *Analytica Chimica Acta*, 1029:50–57, 2018. ISSN 0003-2670. doi: 10.1016/j.aca.2018.05.001.

T.-Y. Lin, P. Goyal, R. Girshick, K. He, and P. Dollár. Focal loss for dense object detection. In *Proceedings of the IEEE international conference on computer vision*, pages 2980–2988, 2017.

W. J. Nash, J. B. Ngere, L. Najdekr, and W. B. Dunn. Characterization of Electrospray Ionization Complexity in Untargeted Metabolomic Studies. *Analytical Chemistry*, 96 (27):10935–10942, July 2024. ISSN 0003-2700. doi: 10.1021/acs.analchem.4c00966.

A. Paszke, S. Gross, F. Massa, A. Lerer, J. Bradbury, G. Chanan, T. Killeen, Z. Lin, N. Gimelshein, L. Antiga, et al. Pytorch: An imperative style, high-performance deep learning library. *Advances in neural information processing systems*, 32, 2019.

A. Roux, Y. Xu, J.-F. Heilier, M.-F. Olivier, E. Ezan, J.-C. Tabet, and C. Junot. Annotation of the Human Adult Urinary Metabolome and Metabolite Identification Using Ultra High Performance Liquid Chromatography Coupled to a Linear Quadrupole Ion Trap-Orbitrap Mass Spectrometer. *Analytical Chemistry*, 84(15):6429–6437, Aug. 2012. ISSN 0003-2700. doi: 10.1021/ac300829f.

S. Samino, M. Vinaixa, M. Díaz, A. Beltran, M. A. Rodríguez, R. Mallol, M. Heras, A. Cabre, L. Garcia, N. Canela, F. de Zegher, X. Correig, L. Ibáñez, and O. Yanes. Metabolomics reveals impaired maturation of HDL particles in adolescents with hyperinsulinaemic androgen excess. *Scientific Reports*, 5(1):11496, June 2015. ISSN 2045-2322.

doi: 10.1038/srep11496.

O. Senan, A. Aguilar-Mogas, M. Navarro, J. Capellades, L. Noon, D. Burks, O. Yanes, R. Guimerà, and M. Sales-Pardo. CliqueMS: A computational tool for annotating in-source metabolite ions from LC-MS untargeted metabolomics data based on a coelution similarity network. *Bioinformatics*, 35(20):4089–4097, Oct. 2019. ISSN 1367-4803, 1460-2059. doi: 10.1093/bioinformatics/btz207.

C. A. Smith, E. J. Want, G. O'Maille, R. Abagyan, and G. Siuzdak. XCMS: Processing mass spectrometry data for metabolite profiling using nonlinear peak alignment, matching, and identification. *Analytical Chemistry*, 78(3):779–787, Feb. 2006. ISSN 0003-2700. doi: 10.1021/ac051437y.

M. Tancik, P. Srinivasan, B. Mildenhall, S. Fridovich-Keil, N. Raghavan, U. Singhal, R. Ramamoorthi, J. Barron, and R. Ng. Fourier features let networks learn high frequency functions in low dimensional domains. *Advances in neural information processing systems*, 33:7537–7547, 2020.

R. Tautenhahn, C. Böttcher, and S. Neumann. Highly sensitive feature detection for high resolution LC/MS. *BMC Bioinformatics*, 9(1):504, Nov. 2008. ISSN 1471-2105. doi: 10.1186/1471-2105-9-504.

K. Uppal, D. I. Walker, and D. P. Jones. xMSannotator: An R Package for Network-Based Annotation of High-Resolution Metabolomics Data. *Analytical Chemistry*, 89(2): 1063–1067, Jan. 2017. ISSN 0003-2700. doi: 10.1021/acs.analchem.6b01214.

P. Veličković, G. Cucurull, A. Casanova, A. Romero, P. Lio, and Y. Bengio. Graph attention networks. *arXiv preprint arXiv:1710.10903*, 2017.

R. Wang, M. Lu, S. An, J. Wang, and C. Yu. 3D-MSNet: A point cloud based deep learning model for untargeted feature detection and quantification in profile LC-HRMS data. *Bioinformatics*, 39(5):btad195, 2023. ISSN 1367-4811. doi: 10.1093/bioinformatics/btad195.
